# Pan-cancer multi-omics analysis identifies non-canonical oncogenic functions beyond DNA replication of the POLE2 subunit of DNA Polymerase epsilon

**DOI:** 10.64898/2025.12.09.693296

**Authors:** Alina Krasilia, Deepali Chaudhry, Iwona J. Fijalkowska, Michal Dmowski

**Author notes:** Corresponding author: A.K.; M.D.

## Abstract

DNA polymerase epsilon (Polε) is essential for high-fidelity leading-strand synthesis. It comprises a catalytic subunit, POLE and three accessory subunits, including POLE2. In contrast to POLE, the biological functions and clinical significance of POLE2 remain unexplored. POLE2 is vital for the assembly of Polε and for interactions with the CMG helicase. Yeast studies show that dysfunction of its ortholog, Dpb2, cause genomic instability and impaired cell cycle progression. To investigate the significance of POLE2, this study provides a comprehensive multi-omics analysis across tumor types. We show that POLE2 upregulation is associated with decreased overall survival in a cancer-type-specific manner. Its expression levels, rather than copy number variation, emerge as a more reliable prognostic marker. POLE2 expression regulatory mechanisms include promoter methylation, isoform usage, and epigenetic regulators. Elevated POLE2 mRNA levels are rarely followed by increased protein levels, possibly due to abundant non-coding isoforms. The localization of mutations in POLE2 in cancer samples suggests an effect on its interaction with the catalytic subunit of Polε. Co-expression and pathway enrichment analyses reveal a connection between POLE2 and transcriptional regulation involving E2F, EZH2, and RB. Our findings highlight POLE2 as a contributor to genome stability, a candidate cancer biomarker, and a possible therapeutic target.

## INTRODUCTION

Faithful DNA replication is essential for preserving genomic integrity, and its disruption is a major contributor to mutagenesis and oncogenesis. DNA polymerase epsilon (Polε) is one of the key enzymes involved in high-fidelity DNA synthesis, primarily responsible for leading-strand replication [1]. It comprises a catalytic subunit (POLE) and three accessory subunits—POLE2, POLE3, and POLE4—involved in the holoenzyme complex assembly and function [2–5]. While the catalytic subunit POLE has been extensively studied, particularly in its proofreading exonuclease domain and its role in hypermutated tumors, the functions and mutational consequences of its non-catalytic subunits remain less understood.

Although it lacks enzymatic activity, POLE2 is required for viability in yeast and mammalian cells [6–8]. Moreover, POLE2 is essential for the assembly and stability of the Polε holoenzyme, i.e., for anchoring the catalytic core to accessory subunits [6–8] and for stabilizing interactions with the replication helicase within the CMG (Cdc45-MCM-GINS) complex [2,5,9]. Structural studies have shown that POLE2 is highly conserved from yeast (Dpb2) to humans, indicating a fundamental role in eukaryotic DNA replication [6,7,10,11]. Dysregulation or mutation of POLE2 may impair the cooperation between Polε and the CMG helicase complex, potentially leading to replication stress, increased mutation rates, and genomic instability—hallmarks of cancer.

So far, our knowledge of the effects of dysfunctions of the main accessory subunit of Polε comes from studies of the yeast’s ortholog of POLE2, *DPB2*. In our previous studies, it has been shown that mutations in *DPB2,* which affect the interactions between the polymerase and helicase complexes, result in strong genomic instability. This involves increased mutation rates, impaired progression through the S phase, abnormal cell morphology, defective replication checkpoint activation, and increased instability of DNA repeat tracts [12–17]. The impaired functioning of the replisome in Polε-defective cells is at the source of perturbations in helicase-polymerase function, severely affecting DNA replication and resulting in the formation of single-stranded DNA (ssDNA). This necessitates a rescue by DNA repair mechanisms, mainly those related to homologous recombination [18]. Moreover, it affects the division of labor between the two main DNA polymerases, resulting in an increased contribution of Pol δ to leading strand replication [19].

Recent reports have identified a range of POLE2 alterations in various tumor types, including colorectal, endometrial, renal, biliary, and breast cancers [20–26]. These alterations include missense and nonsense mutations, frameshift insertions and deletions, silent mutations with potential regulatory impact, and large-scale structural variations. Moreover, changes in POLE2 mRNA expression and epigenetic modifications, such as promoter methylation, have also been documented; however, their functional implications are not yet fully understood. Notably, the recurrence of specific POLE2 mutations across multiple tumor types suggests that selective pressure may favor replication-disruptive variants during tumor evolution.

This study presents a comprehensive analysis of POLE2 alterations and their potential clinical and functional consequences. We evaluate the association between POLE2 status and patient overall survival to determine whether POLE2 alterations may serve as prognostic markers. Additionally, we assess POLE2 gene expression, DNA methylation status, and alternative isoform usage, exploring how these layers of regulation may influence cancer phenotype. We also analyze the spectrum of POLE2 mutations across multiple cancers using publicly available datasets, including the COSMIC database. To understand the broader impact of POLE2 dysregulation, we perform genetic alteration analysis, identify co-expressed genes, and carry out enrichment analysis of associated pathways and biological processes. Through this integrative multi-omics approach, we aim to elucidate the role of POLE2 in cancer biology, its potential contribution to genome instability, and its relevance as a biomarker or therapeutic target. Given the central importance of DNA replication in oncogenesis, a deeper understanding of POLE2 may uncover novel mechanisms of tumor development and progression.

## MATERIALS AND METHODS

### Overall Survival Analysis Based on POLE2 Molecular Alterations

Four independent TCGA-based platforms were employed to assess the association between POLE2 expression and overall survival (OS): UCSC Xena, ENCORI, GEPIA, and TISIDB. Among these, UCSC Xena (University of California, Santa Cruz; https://xena.ucsc.edu/)[27] was used for pan-cancer analysis and further individual cancer-type-specific analysis. In the initial pan-cancer analysis, the Kaplan-Meier Plot option in UCSC Xena was used to evaluate survival associations using POLE2 copy number variation (CNV), average mRNA expression across all cancer types, and mutation status. Following this, a second stage of analysis was performed at the level of individual cancer types. Initially, the Kaplan-Meier Plot option in UCSC Xena was used for each TCGA cancer type (listed in Table S1) to evaluate the significance of the association between POLE2 expression and OS. To validate and reinforce the findings, results were compared against three additional platforms: ENCORI (Encyclopedia of RNA Interactomes; https://rnasysu.com/encori/)[28] via its Pan-Cancer Survival Analysis module to identify cancer types showing statistically significant associations (p < 0.05) between POLE2 expression and OS; GEPIA2 (Gene Expression Profiling Interactive Analysis; http://gepia2.cancer-pku.cn/) [29] using OS analysis with a 50% expression cutoff to define high- and low-expression cohorts; TISIDB (Tumor-Immune System Interactions Database; http://cis.hku.hk/TISIDB/)[30] through its Clinical module for OS assessment across TCGA datasets in relation to POLE2 expression.

TCGA cancer types (listed in Table S1) that showed consistent and significant OS associations with POLE2 expression across all four platforms were selected for focused visualization. For these cancer types, UCSC Xena’s pre-processed survival data—including group stratifications (e.g., high vs. low expression)—were exported and used for survival curve visualization in RStudio [31], employing the *survival* package [32][33]. The Surv() function was used to define survival time and event status. Kaplan-Meier curves were fitted using survfit(), and group differences were evaluated with the log-rank test via survdiff(). P-values were calculated from the resulting chi-squared statistics using pchisq().To quantify the effect of POLE2 expression on overall survival, a Cox proportional hazards regression model was fitted using the coxph() function from the *survival* package in R. POLE2 expression group (“High” vs. “Low”) was used as a categorical explanatory variable. The model estimated the hazard ratio (HR) and its 95% confidence interval (CI).Visualization was performed using the *ggplot2* package [34], and multiple plots were arranged using wrap_plots() from *patchwork* [35] for figure composition. For most of the graphs, theme_few() from package *ggthemes* [36] was used.

### mRNA expression analysis

Average mRNA and Exon expression of POLE2 in cancer and normal TCGA PANCAN dataset were downloaded from UCSC Xena (University of California, Santa Cruz; https://xena.ucsc.edu/). Alongside general expression analysis, POLE2 mRNA levels in different copy number variation groups were analyzed, particularly from the “gistic2_thresholded” dataset, which includes only tumor samples, while only “Primary tumor” sample type was taken into analysis. In this dataset, CNV groups were divided into five categories: -2 represents a homozygous deletion, where both gene copies are likely lost, although some mRNA expression may persist due to normal cell contamination or technical artifacts. -1 indicates a heterozygous deletion, with one copy lost. 0 corresponds to a normal (diploid) copy number. +1 denotes a low-level gain, typically one extra copy, while +2 indicates a high-level amplification involving multiple additional copies. These simplified categories approximate underlying genomic alterations but may not fully capture accurate gene dosage.

The significance of the difference between groups was calculated using RStudio. Pairwise comparisons between cancer and normal sample types were conducted using the Wilcoxon rank sum test. The Benjamini-Hochberg correction was applied to control for false discovery due to multiple comparisons. The analysis was performed using the pairwise.wilcox.test() function from the *rstatix*[37] package. The stat_pvalue_manual() function from ggpubr[38] package was used to visualize significance on the plot.

For cancer-specific expression analysis, the UCSC Xena database was used as a source of expression data, while other databases were used to validate cancer types that consistently show significant differences between cancer and normal samples. GEPIA2 (Gene Expression Profiling Interactive Analysis; http://gepia2.cancer-pku.cn/) [29] database with and without including GTEx normal tissue data was used inside via the “Expression DIY → Box Plot” panel. A P-value cutoff of 0.05 and a Log2FC of 1 were used. GEPIA2 applies a one-way ANOVA model, treating the disease state (Tumor vs. Normal) as the grouping variable. Expression values were log-transformed as log₂(TPM + 1) prior to analysis. Cancer types that resulted in significant changes between cancer and normal were noted. Additionally, **ENCORI** (Encyclopedia of RNA Interactomes; https://rnasysu.com/encori/) [28] Pan-Cancer Differential Expression Analysis option was used to assess TCGA cancer types with a significant difference in POLE2 expression between cancer and normal samples. Similarly, UALCAN (University of Alabama at Birmingham Cancer data analysis Portal, https://ualcan.path.uab.edu/analysis.html)[38,39] was used for the same purpose. To further validate the data, TIMER2.0 (http://timer.comp-genomics.org/) [40,41] database “Cancer Exploration→ Gene_DE”panel was used, and cancer types with p<0.05 were selected.

TCGA data from the UCSC Xena database were used to visualize selected cancer types. Statistical comparisons between cancer and normal samples were performed using the wilcox.test() function from the *rstatix* package. Visualization was performed as described in Section 2.1.

For visualization of POLE2 expression in different tissues, the data was collected using the getTissueVST() function from the *correlationAnalyzeR* package [42]. Statistical comparisons between cancer and normal samples were performed using the wilcox.test() function from the *rstatix* package. Effect sizes were assessed using two metrics: log2 fold change (log2FC) to quantify the magnitude of expression change, and Cohen’s d to measure the standardized mean difference between cancer and normal samples. Cohen’s d were computed using the cohen.d() function from the *effsize*[43] package with a 95% confidence interval. Cohen’s d represents the standardized mean difference between two groups, expressed in units of pooled standard deviation. Visualization was performed as described above.

Gene expression data from TCGA and GTEx were obtained from the UCSC Xena database to compare the expression of genes involved in DNA replication and repair pathways across various TCGA cancer types. GTEx samples were matched to corresponding TCGA cancer types using a manually curated tissue concordance table, based on the “primary_site” metadata provided by UCSC Xena. Differential expression analysis was performed using the limma[45] package for each cancer type. Expression matrices were transposed so that genes were rows and samples were columns. A contrast model comparing cancer to normal samples was fit using the lmFit() and eBayes() functions. All genes were tested, and the log2 fold change (logFC), raw p-value, and adjusted p-value (using the Benjamini–Hochberg method) were calculated for each. To visualize differential expression patterns of genes involved in DNA replication and repair, a heatmap was generated using *`ggplot2`*[34]. Genes were grouped into functional categories (e.g., MCM, GINS, Pol ε, CDC45), and each group was assigned a consistent color. Gene names were displayed in color using markdown formatting via *`ggtext`*[44], and a custom legend indicating gene group membership was added alongside the plot using the *`cowplot`*[45] package.

POLE2 mRNA expression across tumor stages was analyzed using batch effect–normalized RNA-seq expression (RSEM) data from the UCSC Xena database and cBioportal (https://www.cbioportal.org/) [46–48]. In the UCSC Xena TCGA dataset, ajcc_pathologic_tumor_stage was used as the phenotypic stage annotation alongside POLE2 expression data, which were converted from log2(norm + 1) format back to normalized RSEM counts. Tumor stage codes were standardized to a numeric scale from 1 to 4. In cBioPortal, the TCGA PanCancer Atlas Studies were selected as the data source. POLE2 expression values, already provided on a linear scale, were analyzed against the clinical attributes of the American Joint Committee on Cancer Tumor Stage Code. Starting from this point, both datasets were processed identically. Outliers were removed within each cancer type and stage using the interquartile range (IQR) method. Cancer types with significant stage-specific differences in POLE2 expression were identified using the Kruskal-Wallis test, implemented via the kruskal_test() function from the rstatix package. For these cancer types, pairwise Wilcoxon rank-sum tests (pairwise_wilcox_test()) were performed to compare expression levels between stages, with p-values adjusted using the Benjamini-Hochberg method.

For protein and mRNA expression comparison, UALCAN portal was used to export CPTAC plot data for cancer types comparable with TCGA cancers, together with TCGA plot data for the same cancer types. Visualization was performed as described in Section 2.1.

### POLE2 methylation analysis

DNA methylation for POLE2 promoter region from Illumina HumanMethylation450 BeadChip (450K array) dataset was downloaded from the UCSC Xena (University of California, Santa Cruz; https://xena.ucsc.edu/)[27] database. Gene average methylation and detailed view data were exported together with cancer type abbreviation and sample type data. Methylation levels (β-values) between cancer and normal samples were compared using the Wilcoxon rank-sum test, implemented via the wilcox.test() function from the *rstatix* package. For comparison between CpG sites and cancer types, p-values were adjusted using the Benjamini-Hochberg method. The mean methylation difference (Δβ) was calculated as the difference in average β-values between cancer and normal groups. Visualization was performed as described in Section 2.1.

The location of reported CpG sites relative to the POLE2 sequence was exported from UCSC Genome Browser (https://genome.ucsc.edu/)[49]. The Human (GRCh37/hg19) genome assembly was used for analysis. CpG site coordinates were obtained from the “Illumina 450K Methylation Array” subtrack under the “Array Probesets” track [50]. CpG island locations were retrieved from the “CpG Islands” track [51]. Gene annotation for POLE2 was based on the “GENCODE V47lift37” track [52], and functional domain information, including the N-terminal domain of POLE2, was obtained from the “Pfam in UCSC Gene” track [53].

### Isoform expression and transcript analysis

TOIL RSEM isoform percentage data from the UCSC Xena database was used for isoform expression analysis. Isoform percentage and sample type data were exported from the TCGA and GTEx datasets. GTEx normal tissue data were assigned to corresponding TCGA cancer type abbreviations for comparison, as detailed in Table S2. For the Cancer category, only samples labeled as “Primary Tumor” from TCGA were used. The Normal_TCGA category included “Solid Tissue Normal” samples, representing tissue adjacent to tumors. The Normal_GTEx category included “Normal” samples from GTEx, representing normal tissues unrelated to tumors. Isoform structure data were exported from the UCSC Genome Browser, using the Comprehensive Gene Annotation Set from GENCODE Version 29 (Ensembl 94) based on the human genome assembly GRCh38/hg38. Transcripts were classified based on their annotated biological function as coding (protein-coding and polymorphic pseudogenes), non-coding (non-protein-coding transcripts), or problematic (biotypes including retained_intron, TEC, or disrupted_domain) [54]. Isoform expression levels for each TCGA cancer type were exported from GEPIA 2.0.

### Genetic alteration analysis

A systematic analysis of POLE2 mutations across cancer types was conducted using data from the Catalogue of Somatic Mutations in Cancer (COSMIC)(https://www.cosmickb.org/) and the UCSC Genome Browser (Human GRCh38/hg38) (https://genome.ucsc.edu/) [49]. The POLE2 gene was first aligned with the COSMIC mutation track (Version 101) [55] in the UCSC Genome Browser to extract mutation coordinates. Descriptive data were then obtained from the COSMIC database for the ENST00000216367.9 transcript (POLE2-001), including information on point mutation positions, amino acid changes, and associated tissue types (Table S5). Exon-intron coordinates were obtained from the UCSC Genome Browser for the same transcript. Somatic mutations with a recurrence count greater than one were visualized using ggplot2 [34] and cowplot [45] packages in R.

### Correlation analysis

Lists of genes with high POLE Pearson correlation value (>0.8) were exported from 4 independent databases, where Correlation AnalyzeR (https://gccri.bishop-lab.uthscsa.edu/shiny/correlation-analyzer/) [42] contained data both for cancer and normal samples, UALCAN (University of Alabama at Birmingham Cancer data analysis Portal, https://ualcan.path.uab.edu/analysis.html)[38,39] and GEPIA2 (Gene Expression Profiling Interactive Analysis; http://gepia2.cancer-pku.cn/) [29] provided data only for cancer patients, GTExPortal (https://gtexportal.org/home/) – exclusively for normal tissue. Initial data sources were TCGA data for UALCAN and GEPIA2, ARCHS4 for Analyzer, and GTEx original data for GTExPortal. In the Correlation AnalyzeR database, POLE2 correlated genes were exported for available tissue types with both cancer and normal sample data. The UALCAN database provides POLE2-correlated genes for TCGA cancer samples through the “Correlation” section. The GEPIA2 database offers the top 1000 genes most similar to POLE2 in TCGA tumor samples, available under the “Expression Analysis → Similar Genes Detection” tab. To get POLE2 correlation values for normal tissues, open-access data from GTExPortal was downloaded, namely GTEx Analysis V10 RNA-seq data (GTEx_Analysis_2022-06-06_v10_RNASeQCv2.4.2_gene_tpm_non_lcm.gct) with metadata (GTEx_Analysis_v10_Annotations_SampleAttributesDS.txt). Sample IDs were extracted for each tissue, and gene expression data were filtered to include only relevant samples (Table S3).

Correlation between POLE2 expression and all other genes was computed using Pearson correlation for each tissue using the cor.test() function. P-values for the correlations were calculated, and genes with p < 0.05 and |correlation| > 0.8 were retained. Tissues were grouped into functional categories based on anatomical and biological relevance using a predefined mapping table (Table S4).

Gene biotypes were determined from the GENCODE v47 Basic Annotation GTF file. The GTF file was parsed to extract gene_name and gene_type, retaining unique entries. Genes annotated as “protein-coding” were selected for downstream analysis.

To visualize the overlap between POLE2-correlated or similar genes obtained from multiple datasets (UALCAN, GEPIA2, GTEx, and Analyzer), the UpSet plot was generated using the upset() function from the ComplexUpset package[56], with ‘exclusive_intersection’ mode. Intersections containing at least one gene were extracted using the upset_data()function from the ComplexUpset package and were used for subsequent enrichment analysis.

### Enrichment analysis

Over-representation analysis (ORA) was performed to identify pathways enriched among genes uniquely shared across dataset intersections. Genes from each exclusive intersection were tested against predefined gene sets (e.g., Hallmark pathways) using the enricher() function from the clusterProfiler package [57] with a false discovery rate (FDR) threshold (q-value cutoff) of 0.2. P-values were adjusted using the Benjamini–Hochberg method to control for multiple testing. Gene set annotations were loaded using read.gmt() from clusterProfiler. Significant results (adjusted p-value < 0.05) were visualized as dot plots with ggplot2 as described in Section 3.1. Predefined gene sets in .gmt format were downloaded from MSigDB (Molecular Signatures Database)( https://www.gsea-msigdb.org/gsea/msigdb)[58], specifically from “Human MSigDB v2024.1.Hs” updated August 2024. In Fig. 8, the “h.all.v2024.1.Hs.symbols.gmt” [59] file was used, whereas in Fig. 9B, the “h.all.v2024.1.Hs.symbols.gmt”, “c2.cp.kegg_legacy.v2024.1.Hs.symbols.gmt”, “c6.all.v2024.1.Hs.symbols.gmt”, and “c3.tft.tft_legacy.v2024.1.Hs.symbols.gmt”[59,60] files were used. Enriched terms and their genes were visualized with the *plotly* package [61] and saved with the *webshot2* package [62].

Functional enrichment analysis was performed using the Metascape (http://metascape.org) [63] resource based on exclusively cancer-related gene sets for separate TCGA cancer types. Circos plot visualization and protein-protein interaction network (PPI network, Physical Core option) was generated by Metascape, using Molecular Complex Detection (MCODE) algorithm, based on data from STRING, BioGrid, OmniPath, and InWeb_IM. PPI network was edited in Cytoscape[64].

## RESULTS AND DISCUSSION

### Survival analysis reveals the prognostic significance of POLE2 expression in cancer

To assess how POLE2 perturbations affect the overall survival of cancer patients, Kaplan–Meier (KM) plots were generated for POLE2 CNV, mRNA expression, and mutations (Fig. 1A). The results show that overall survival was significantly lower in patients with higher POLE2 mRNA expression (high expression group: n = 4828; low expression group: n = 4814; p-value < 2.2e-16). In contrast, no significant differences in survival rates were observed between groups stratified by POLE2 copy number variation (CNV) (high CNV group: n = 4931; low CNV group: n = 4883; p-value = 0.5418) or mutation status (mutation group: n = 59; no mutation group: n = 8627; p-value = 0.6421). Notably, POLE2 mutations were rare across the cohort, and the small number of mutated cases compared to the non-mutated group makes it difficult to draw definitive conclusions about the impact of mutations on survival.

**Figure 1.**
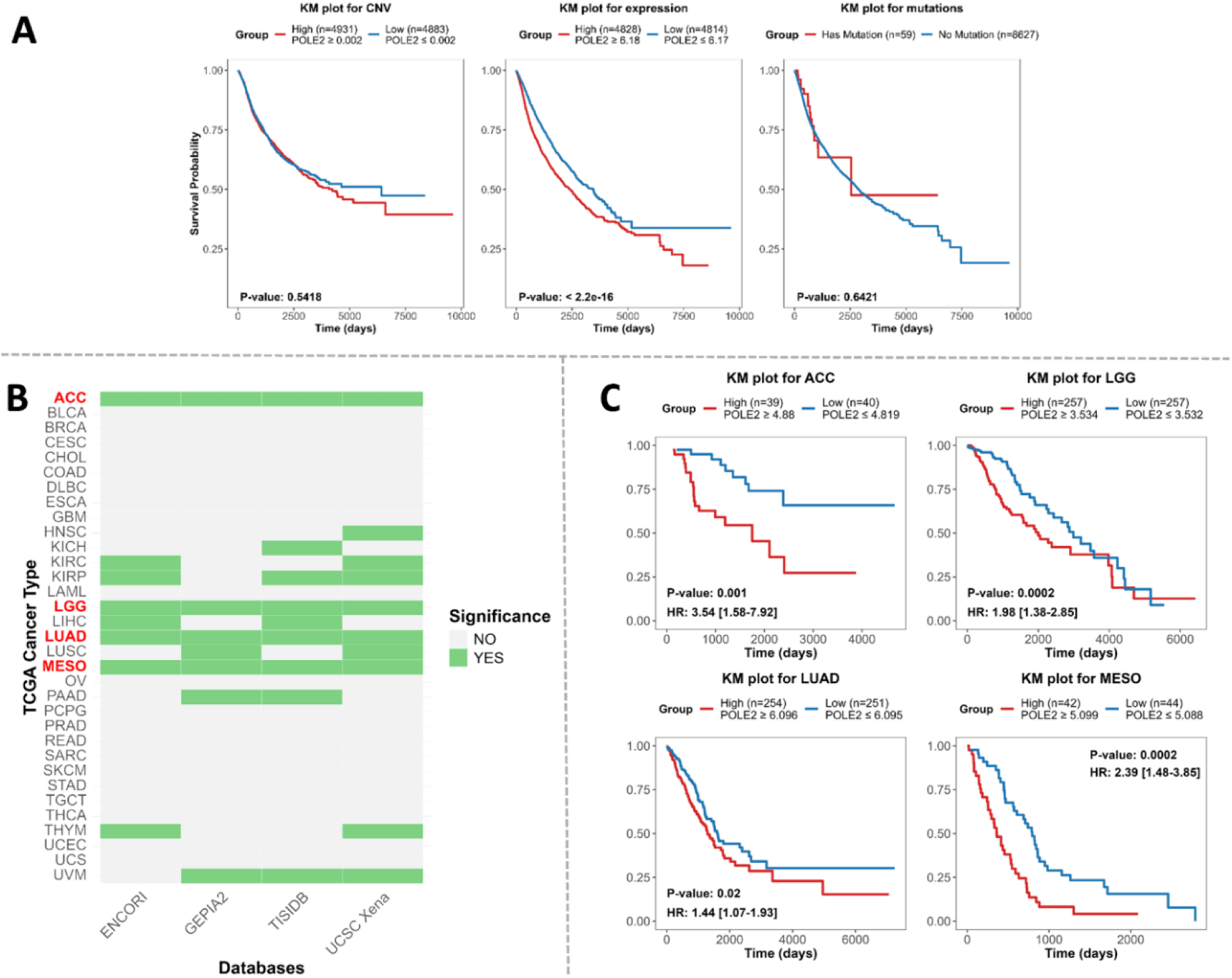
Overall survival analysis of patients based on POLE2 alterations. **A.** Kaplan–Meier survival plots from TCGA Pan-Cancer (PANCAN) data from the UCSC Xena database assessing the impact of POLE2 copy number variation (CNV), mRNA expression levels, and mutation status on patient overall survival. **B.** Heatmap summarizing the statistical significance (*p* ≤ 0.05, green) of POLE2 mRNA expression-associated survival across multiple TCGA cancer types, evaluated using independent databases (ENCORI, TISIDB, GEPIA2, UCSC Xena). Cancer types consistently showing significant associations across all platforms are highlighted in red. **C.** Kaplan–Meier plots from TCGA Pan-Cancer (PANCAN) data from the UCSC Xena database for selected cancer types from panel B (ACC, LGG, LUAD, MESO) in which high POLE2 mRNA level consistently correlates with overall survival outcomes across all datasets. The plot displays the hazard ratio (HR) followed by its 95% confidence interval in square brackets. For example, “HR: 3.54 [1.58–7.92]” indicates that patients with high POLE2 expression had a 3.54 times higher risk of death compared to those with low expression, with the true value likely falling between 1.58 and 7.92.

These findings suggest that POLE2 expression level, rather than CNV or mutation status, may serve as a potential prognostic marker for cancer patients. However, CNV and mutation may still exert indirect effects by influencing gene expression or through non-survival-related mechanisms. These aspects have been further explored in subsequent sections of the analysis, where we examine the relationship between CNV and POLE2 expression and the frequency and potential functional impact of POLE2 mutations across cancer types.

To identify particular cancer types in which survival is significantly associated with POLE2 mRNA level, data from 33 TCGA cancer types (listed in Table S1) were collected from 4 independent databases (UCSC Xena, ENCORI, GEPIA, TISIDB) and compared (Fig. 1B). Patients were divided into high and low POLE2 expression groups based on cancer-type-specific thresholds. The analysis revealed four cancer types—ACC, LGG, LUAD, and MESO—consistently showing a significant (p ≤ 0.05) survival difference between groups stratified by POLE2 mRNA level. Separate Kaplan–Meier (KM) plots for these cancer types are presented in Fig. 1C. Corresponding p-values and hazard ratios (HR) with 95% confidence intervals (CI) from Cox proportional hazards models are shown on each plot. The HR indicates the relative risk of death in the high-expression group compared to the low-expression group. The strongest effect was observed in ACC, where high POLE2 expression was associated with significantly poorer survival (HR: 3.54 [1.58–7.92], p = 0.001). MESO also showed a strong association (HR: 2.39 [1.48–3.85], p = 0.0002), followed by LGG (HR: 1.98 [1.38–2.85], p = 0.0002). A weaker but still significant association was found in LUAD (HR: 1.44 [1.07–1.93], p = 0.02).

Several other cancer types also demonstrated partial evidence linking POLE2 mRNA levels to survival outcomes. Specifically, KIRP and UVM showed supportive results in 3 out of 4 databases; KIRC, LIHC, LUSC, PAAD, and THYM showed positive associations in 2 out of 4 databases; HNSC and KICH showed associations in 1 out of 4 databases. Separate Kaplan–Meier (KM) plots for these cancers can be found in Fig. S1. Notably, KICH and UVM exhibited strong hazard ratios (HR: 3.44 [0.71–16.56], HR: 2.95 [1.16–7.51], respectively), although only UVM reached statistical significance, while the wide CI in KICH reflects uncertainty due to a small sample size. A similarly elevated HR was seen in KIRP (1.96 [1.05–3.66]), where the association was statistically significant, indicating a nearly twofold increase in risk. Moderate but significant HRs around 1.4 were found in KIRC and LIHC, indicating a weaker yet consistent increase in risk with higher POLE2 expression.

In contrast, THYM demonstrated an HR well below 1 (0.18 [0.30–0.89]), suggesting a protective effect of high POLE2 expression, which was also statistically significant. LUSC showed a similar inverse trend (HR: 0.75 [0.57–0.99]), though the effect size was smaller. For PAAD and HNSC, HRs close to 1 (1.35 and 0.81, respectively) indicated little to no association, and these findings were not statistically significant.

These findings demonstrate that elevated POLE2 expression is associated with increased hazard ratios and decreased overall survival, with the strength and direction of this association being cancer-type specific. The most pronounced and consistently significant effects were observed in ACC, LGG, LUAD, and MESO. Further examination of Kaplan–Meier curves and hazard ratios in cancer types showing significance in at least one external database indicated that UVM may also have prognostic relevance.

Additionally, KIRC and LIHC displayed hazard ratios comparable to those of LUAD, suggesting similar risk patterns linked to high POLE2 expression. This observation aligns with previous findings in renal cell carcinoma (RCC), of which KIRC is the most common subtype, where high POLE2 expression has been associated with reduced overall and progression-free survival [21]. Notably, although high POLE2 expression has previously been reported to correlate with poor survival in BLCA [26], our data did not support a significant association in this cancer type. In contrast, THYM and LUSC showed an inverse relationship, where elevated POLE2 expression was associated with improved survival, indicating a potential protective effect.

The strategy of cross-validating results across multiple independent databases proved effective, identifying three of the four cancer types with the strongest effects and reinforcing the observed associations’ reliability.

### POLE2 expression analysis in cancer and normal samples

To explore how different alterations of the POLE2 gene expression differ between cancer and normal tissues, we first analyzed POLE2 mRNA expression levels and exon-level expression profiles using data aggregated across all TCGA cancer types. The statistical comparison of mRNA level and Exon expression of the POLE2 gene revealed that there is a significant difference between cancer and normal samples, with an increase in cancer cells (Fig. 2A). To investigate whether specific copy-number variations contribute to the increase in POLE2 expression, the POLE2 mRNA expression analysis was performed across different CNV categories demonstrating significant differences (Fig. 2B). Tumors with 1-copy deletion (n = 1968) showed significantly lower expression compared to tumors with amplification (n = 1220) and high amplification (n = 66). Similarly, tumors with no CNV changes (n = 6043) exhibited significantly lower expression than those with amplification and high amplification. A significant difference was also detected between the amplification and high amplification groups. In contrast, no significant differences were observed between tumors with 2-copy deletions (n = 6) and other categories, likely due to the very small sample size. However, since tumors with one-copy deletion and tumors without CNV changes did not show significant differences in POLE2 expression, these results suggest that while CNV amplification is associated with increased POLE2 expression, copy-number loss or stability alone does not fully account for POLE2 expression variability.

**Figure 2.**
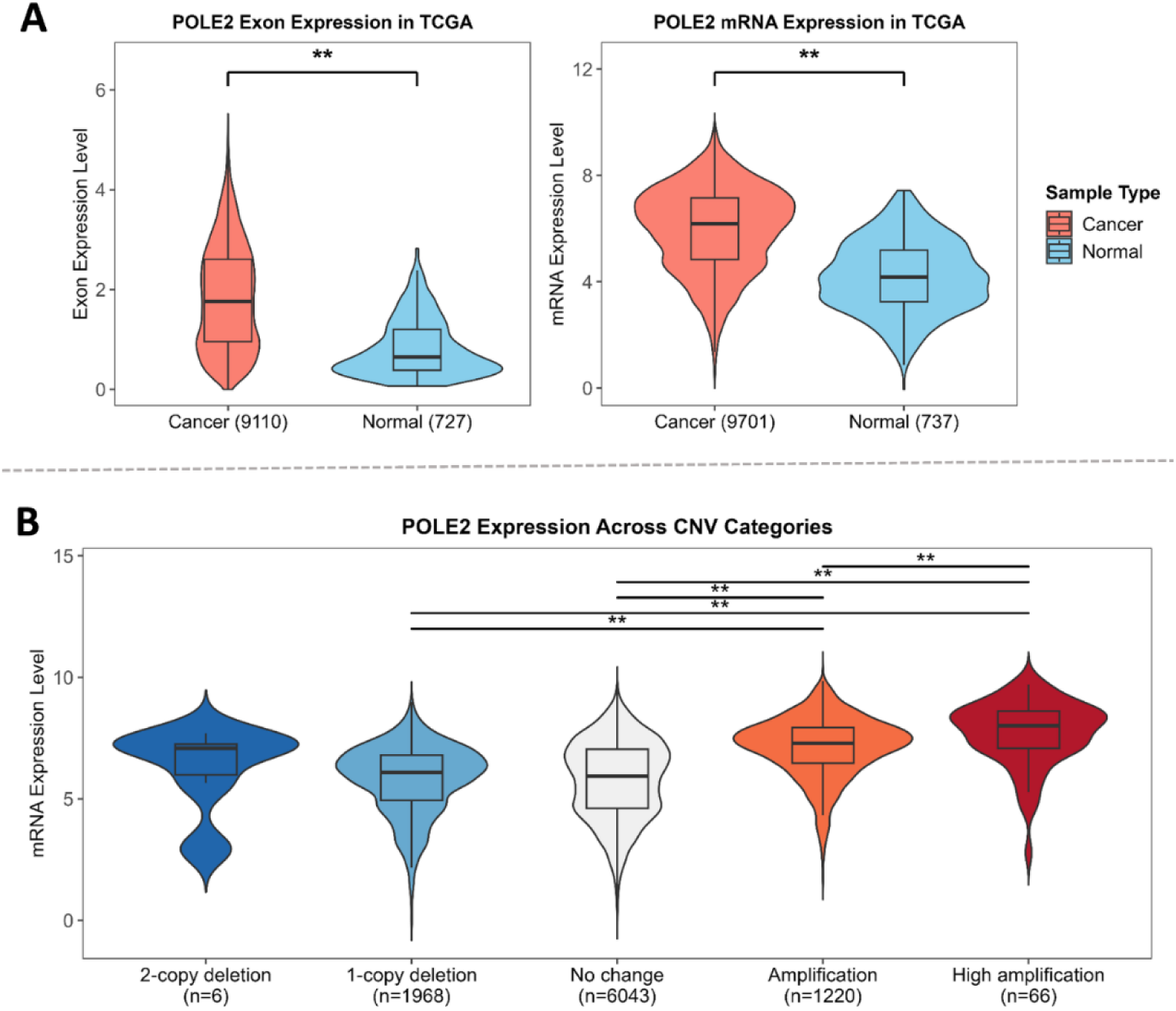
POLE2 expression in cancer and normal tissues based on TCGA data. **A.** POLE2 exon-level expression (left) and POLE2 mRNA expression levels across TCGA cancer samples and normal tissue samples. Expression distributions are shown as violin plots with embedded boxplots. **B.** POLE2 mRNA expression levels across CNV(copy-number variations) categories from TCGA data, where “Amplification” means 1-copy gain and “High amplification” means two or more copy gain. Statistical significance was assessed using the Wilcoxon rank sum test (Mann–Whitney U test). Significance thresholds: p ≤ 0.01 (**), p ≤ 0.05 (*).

To further investigate the relationship between POLE2 expression and copy number variation (CNV), a correlation analysis was performed separately for cancer and normal samples (Fig. S3). A weak but statistically significant positive correlation was observed in both groups, with a Pearson correlation coefficient of r = 0.22 (p = 1.48e-102) in cancer samples and r = 0.25 (p = 2.28e-10) in normal samples (Fig. S3). These findings support that while CNV contributes to POLE2 expression levels, additional regulatory factors are likely to be involved.

To investigate POLE2 expression changes in individual cancer types, we collected data on significant expression differences between cancer and normal tissues from several public databases, including UCSC Xena, GEPIA2 (with and without GTEx), ENCORI, UALCAN, and TIMER. A heatmap summarizing statistically significant differences in POLE2 mRNA expression between cancer and normal samples for each cancer type, as reported by individual databases is shown in Fig. 3A. The comparison revealed that eight cancer types — BLCA, BRCA, ESCA, HNSC, LUAD, LUSC, STAD, and UCEC — showed consistent significant overexpression of POLE2 in all six databases. Additionally, CESC, COAD, and LIHC demonstrated associations in 5 out of 6 databases; KICH, KIRC, KIRP, READ, and THCA showed associations in 4 out of 6 databases; GBM, CHOL, PAAD, and PRAD showed associations in 3 out of 6 databases. For the eight cancer types where POLE2 overexpression was consistently confirmed across all databases, the expression levels in cancer versus normal tissues are visualized in Fig. 3B. In all cases, POLE2 expression was significantly higher in cancer tissues compared to corresponding normal tissues. To evaluate the magnitude of expression differences between cancer and normal tissues, Cohen’s d effect sizes were calculated for each cancer type. All comparisons showed large effect sizes (d > 1.4), indicating substantial separation between groups. The strongest effects were observed in LUSC (d = 3.203), ESCA (d = 2.866), and UCEC (d = 2.378), suggesting highly consistent and pronounced overexpression of POLE2 in cancer samples relative to normal tissue. LUAD (d = 2.114), BLCA (d = 2.072), and BRCA (d = 1.897) also showed strong effects, while STAD (d = 1.763) and HNSC (d = 1.419), though slightly lower, still met the threshold for a large effect size, which is d > 0.8.

**Figure 3.**
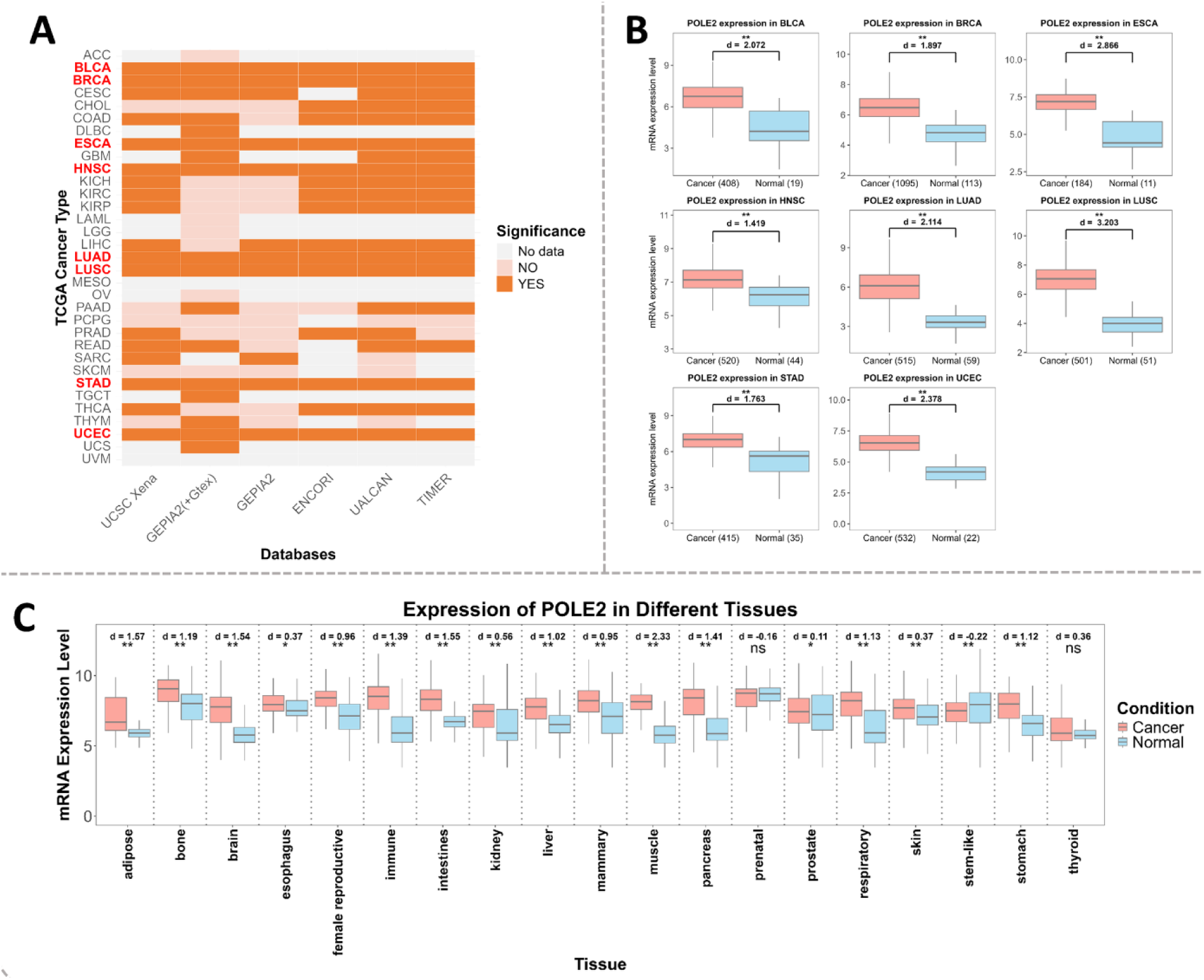
POLE2 mRNA expression in cancer versus normal tissues across databases and tissue types. **A.** Heatmap summarizing the statistical significance of POLE2 mRNA level differences between cancer and normal samples across multiple independent databases, including UCSC Xena, GEPIA2, GEPIA (+GTEx), ENCORI, UALCAN, and TIMER. Cancer types highlighted in red consistently showed significant differences across all platforms. **B.** Boxplots of POLE2 expression based on TCGA RNA-seq data for selected cancer types with consistent significance across databases shown in panel A. Each plot displays the standardized mean difference (Cohen’s *d*). **C.** POLE2 mRNA expression levels across different tissues from ARCHS4 human RNA-seq data, comparing cancer (salmon) and normal (skyblue) samples.

Additional cancers with significant POLE2 expression differences in at least three databases are shown in Fig. S2. The most pronounced effect was observed in CESC (d = 5.421), though this finding is based on only three normal samples and should be interpreted cautiously. Notably strong group separation was also present in KIRP (d = 1.851), LIHC (d = 1.437), KIRC (d = 1.497), and COAD (d = 1.397), all showing large standardized effect sizes. Cancers such as KICH (d = 1.098), PAAD (d = 1.046), READ (d = 1.045), and THCA (d = 0.945) also demonstrated consistent overexpression of POLE2, while PRAD showed a moderate effect (d = 0.554). These findings underscore the consistent and often substantial upregulation of POLE2 in a broad range of tumor types. Consistent with our results, previous studies have reported POLE2 upregulation in specific cancers such as gastric cancer[25]—corresponding well with our STAD findings—as well as in bladder cancer (BLCA) [26], renal cell carcinoma (including KIRC) [21], and osteosarcoma (OS) [65]. These external data further strengthen the notion that POLE2 may play a broader oncogenic role across diverse tumor types and justify continued investigation into its potential biological functions and clinical relevance.

To investigate POLE2 expression patterns beyond TCGA datasets, tissue data from Expression AnalyzeR, which is based on transcriptomic data from the ARCHS4 database, was utilized. This dataset includes a broader range of samples, covering both cancer and normal tissues, including samples derived from cell lines. The expression levels of POLE2 across different tissue types are visualized in Fig. 3C. A statistically significant difference between cancer and normal tissues was observed for nearly all tissue types, except for prenatal and thyroid tissue. The most substantial difference was observed in muscle tissue (d = 2.33), followed by adipose (d = 1.57), intestines (d = 1.55), brain (d = 1.54), pancreas (d = 1.41), immune (d = 1.39), bone (d = 1.19), respiratory (d = 1.13), stomach (d = 1.12), and liver (d = 1.02), all showing large and statistically significant effect sizes (p < 0.01 or p < 0.05). Moderate but significant differences were also seen in female reproductive (d = 0.96), mammary (d = 0.95), esophagus (d = 0.37), skin (d = 0.37), thyroid (d = 0.36), and prostate (d = 0.11). In contrast, prenatal tissue showed a small but significant decrease in POLE2 expression in cancer (d = –0.16), while stem-like tissue showed a similar but non-significant reduction (d = –0.22). This inverse trend may reflect the naturally high POLE2 expression in normal proliferative or developmentally plastic tissues, such as embryonic and stem-like cell populations, where POLE2 may play essential roles in replication and maintenance. In such contexts, malignant transformation might not further elevate its expression, or could even reduce it, depending on cellular differentiation status. These findings indicate that while POLE2 is broadly upregulated in cancer across tissue types, the magnitude and direction of change are tissue-dependent and may be influenced by the intrinsic biology of the tissue of origin. Taken together, TCGA and broader ARCHS4 data highlight specific cancer types and tissues where POLE2 upregulation is consistently observed. These patterns suggest that increased POLE2 expression may act as a tissue-specific indicator of malignancy in select cancer types.

### Differential expression analysis of genes involved in DNA replication and repair pathways

Expression data for a curated set of genes were extracted from the UCSC Xena database to investigate whether specific DNA replication and repair genes are upregulated in particular TCGA cancer types (Fig. 4). These included B-family polymerases such as subunits of polymerase epsilon (POLE, POLE2– 4), polymerase delta (POLD1–4), and polymerase alpha with its primase partners (POLA1, POLA2, PRIM1, PRIM2), which together drive leading and lagging strand synthesis during DNA replication [66]. Additional groups included the X-family polymerases (POLB – Pol β, POLM – Pol μ, POLL – Pol λ), which are specialized for base excision repair (POLB) and non-homologous end joining (POLM, POLL) [67]; the Y-family polymerases (POLH – Pol η, POLI – Pol ι, POLK – Pol κ), which perform translesion synthesis to bypass DNA lesions [68]; and the A-family polymerases (POLQ -Pol θ, POLN -Pol ν, POLG -Pol γ), which participate in mitochondrial DNA replication (POLG), translesion synthesis and base excision repair (POLQ), and error-prone bypass of DNA lesions (POLN) [69]. Only the genes encoding catalytic subunits were included in the analysis for the X-, Y-, and A-family polymerases. Across 20 cancer types, POLE2 stands out as the most overexpressed subunit within the group of replicative polymerases. In contrast, most translesion synthesis and alternative polymerases are downregulated across cancer types. This includes REV3L, POLM, POLL, POLI, POLK, REV1, POLN, and POLG. An exception to this trend is POLQ, which is markedly upregulated despite being a non-replicative polymerase. The consistent upregulation of POLQ presented here aligns well with prior findings [70,71], reinforcing its emerging role in promoting replication stress tolerance and genomic instability in cancer. Notably, TGCT is the only cancer type showing downregulation of POLQ, and it displays particularly strong suppression of POLN, POLI, and POLK. This expression pattern aligns with prior reports indicating that TGCT cells are deficient in key DNA repair pathways, particularly mismatch repair (MMR)[72] and nucleotide excision repair (NER), including interstrand crosslink (ICL) repair[73].

**Figure 4.**
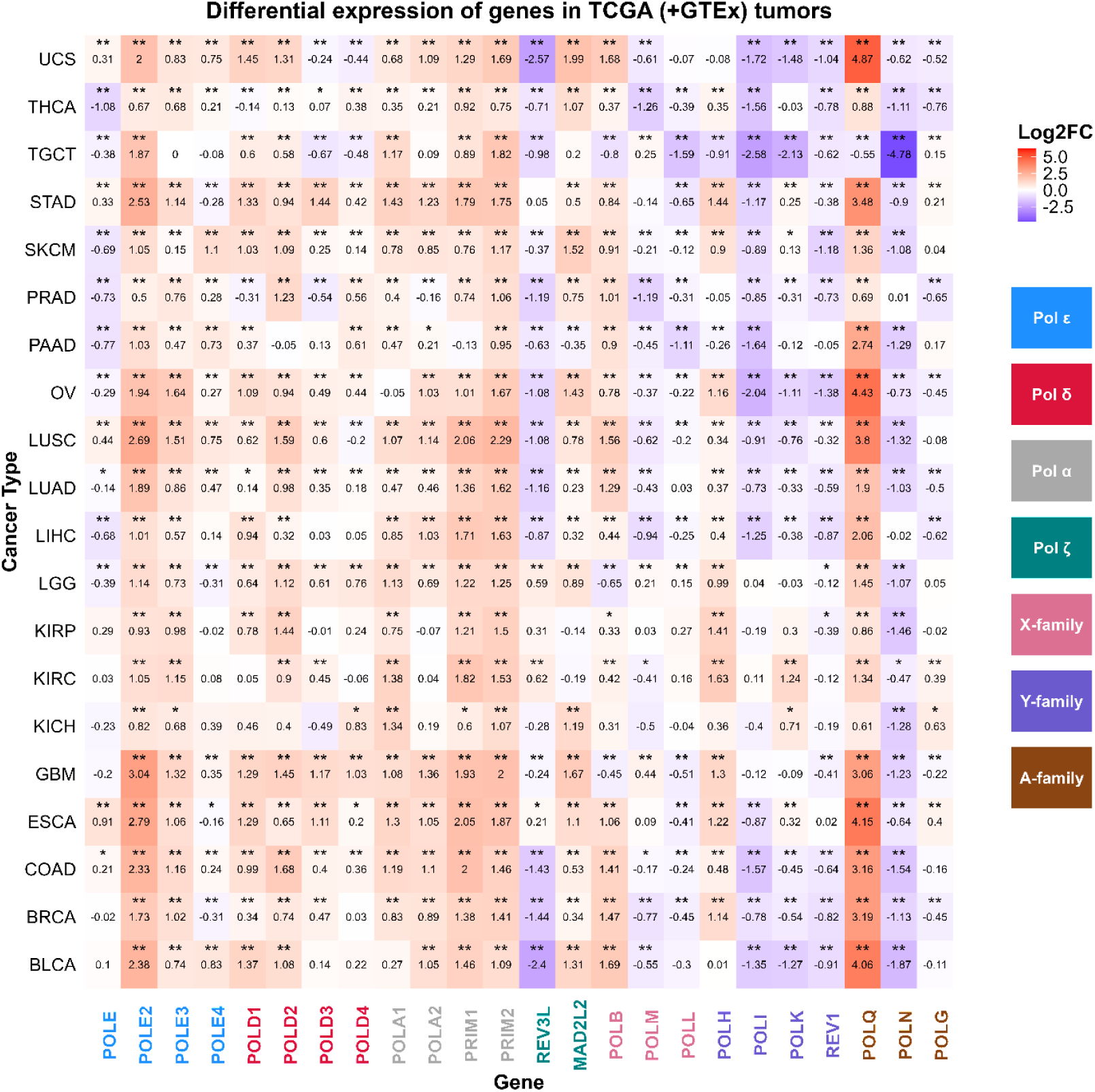
Differential gene expression analysis across selected DNA replication and repair genes in TCGA tumors.

Our findings are consistent with earlier reports showing comparable gene expression profiles. CDC45, for instance, has been shown to be markedly upregulated in various cancers, including thyroid cancer [74], colorectal cancer [75], non-small cell lung cancer [76], esophageal squamous cell carcinoma [77], hepatocellular carcinoma [78], and kidney renal clear cell carcinoma [79].

Likewise, MCM subunits, particularly MCM2, MCM4, MCM5, and MCM7, have been repeatedly reported as overexpressed across diverse tumor types such as breast, lung, gastric, colorectal, ovarian, oral squamous cell carcinoma, and hepatocellular carcinoma [80,81]. These findings have been validated across multiple ethnic cohorts and sample sizes, further supporting the widespread dysregulation of helicase components in tumorigenesis.

The GINS complex, essential for helicase activation, also shows frequent overexpression, with GINS1, GINS2, and GINS4 notably upregulated in sarcoma [82,83], glioma [84], breast [85], and thyroid cancer [86].

Experimental data on the expression of TLS and alternative polymerases remain limited. Contrary to our findings, POLL was reported to be upregulated across multiple cancer types in the study by Starcevic et al. [87]. Notably, that study also documented increased expression of replicative polymerases, including POLA, POLD, and POLE, and overexpression of POLB—findings that align with our observations. Furthermore, it reported predominant underexpression of POLI and POLK, which is consistent with our results.

There is abundant experimental evidence in the literature regarding the overexpression of POLE2 in most tumors, as previously mentioned in this study. Overall, POLE2 emerges as the most consistently upregulated polymerase subunit across 20 TCGA cancer types. The reason for POLE2 upregulation remains unclear. As it lacks catalytic activity and other polymerase subunits are not similarly overexpressed, its role in cancer may not be limited to DNA replication. Components of the replicative helicase, including CDC45, MCMs, and GINS subunits, also show recurrent overexpression. In contrast, translesion and several non-replicative polymerases are broadly downregulated in tumors, except for POLQ, whose strong upregulation suggests a specialized role in the replication stress response.

### POLE2 expression association with tumor progression

To investigate whether POLE2 expression in cancer tissues varies across cancer stages, POLE2 expression data from TCGA were obtained through the UCSC Xena platform and cBioPortal. Among 21 cancer types from UCSC data, only four—namely ACC (P = 0.0154), ESCA (P = 0.0388), LUSC (P = 0.0014), and TGCT (P = 0.0102)—exhibited significant changes in POLE2 mRNA levels among stages (Fig. 5A). In the cBioPortal dataset, the Non-Small Cell Lung Cancer category comprises LUAD and LUSC, while Renal Non-Clear Cell Carcinoma represents two types of TCGA cancers: KIRP and KICH. Statistical analysis of data for 19 cancer groups from cBioPortal identified seven types with significant stage-related variation: ACC (P = 0.00286), KIRP/KICH (P = 0.0349), KIRC (P = 0.00187), BRCA (P = 0.000856), LUAD/LUSC (P = 4.64 × 10⁻⁶), TGCT (P = 0.00351), and CESC (P = 0.00458) (Fig. 5B). To determine more specifically which stages have significantly different POLE2 expression, a pairwise comparison between stages was performed (Fig. 5).

**Figure 5.**
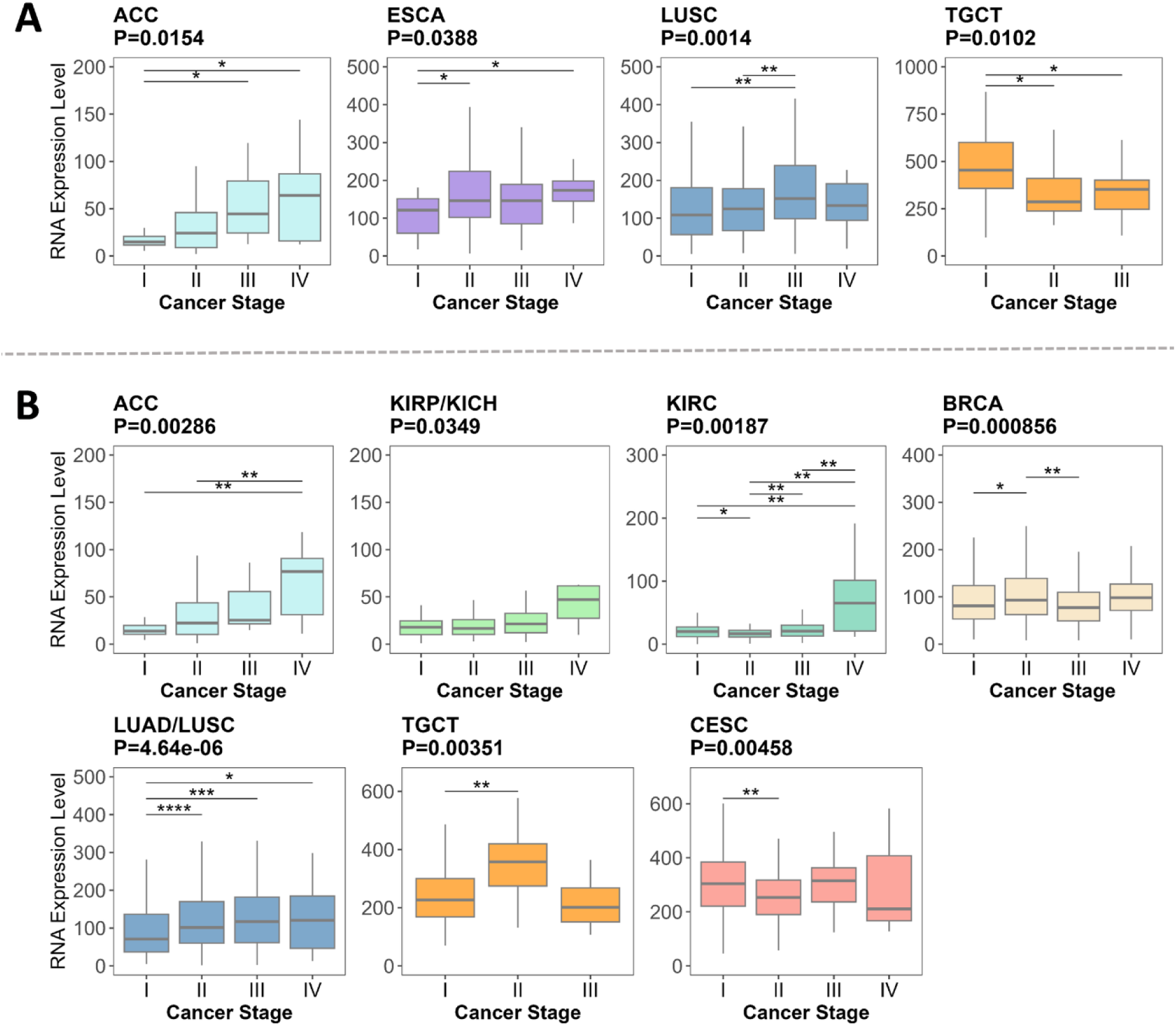
POLE2 mRNA expression levels across cancer stages in selected tumor types. **A.** Batch effect–normalized RNA-seq expression (RSEM) data from the UCSC Xena platform. **B.** Batch effect–normalized RNA-seq expression (RSEM) data from cBioPortal. Cancers shown exhibited statistically significant differences in POLE2 expression across clinical stages. Boxplots display RNA expression levels across stages I–IV. Overall significance was assessed using the Kruskal–Wallis test, followed by pairwise Wilcoxon rank sum tests for post hoc comparisons. P-values were adjusted using the **Benjamini–Hochberg method** to control for multiple testing. Significance thresholds: p ≤ 0.01 (**), p ≤ 0.05 (*).

ACC, KIRP, and KIRC display a clear upward trend in POLE2 expression across cancer stages. In ACC, expression increases significantly between stages I and III, and stages I and IV, with a stronger confirmation of this trend in the second dataset. KIRC shows consistent and pronounced increases in expression, with significant differences observed between almost all adjacent stages, highlighting a progressive upregulation as the cancer advances. KIRP demonstrates a milder upward trend, with statistically significant differences overall, although changes between individual stages are less pronounced. Previous work based on larger renal-cell-carcinoma cohorts has already shown that POLE2 expression rises with tumor progression [21]. However, the role of POLE2 expression in ACC progression was not precisely highlighted. Nevertheless, provided that this study previously showed that POLE2 expression consistently correlates with survival, the influence on POLE2 in ACC needs further investigation.

ESCA and LUAD/LUSC exhibit a slight increase in POLE2 expression as cancer progresses. In ESCA, significant differences are observed between stages I and II, and between stages I and IV, suggesting a modest rise. LUAD/LUSC demonstrates a more apparent stepwise increase, particularly at later stages, with significant differences observed between stage I and all subsequent stages.

LUSC, BRCA, TGCT, and CESC show no strong dependency on POLE2 expression in the cancer stage. LUSC shows a gradual increase in POLE2 expression up to stage III, followed by a slight but non-significant decrease at stage IV. BRCA similarly shows changes between early and later stages, specifically between stages I and II, and I and IV, but overall, the progression is modest. In TGCT from the UCSC Xena database, I stage displays the highest level of POLE2, while in data from cBioPortal, it is true for the II stage, indicating controversial results. CESC shows a difference between stages only between stages I and II, which is insufficient to conclude a clear trend.

Overall, POLE2 expression shows cancer-type-specific stage dependency. In cancers such as ACC, KIRC, and KIRP, expression tends to increase progressively with advancing stage, suggesting a potential link to tumor progression. ESCA and LUAD/LUSC exhibit a modest but noticeable increase in expression across stages. In contrast, cancers like TGCT display an apparent decrease in POLE2 levels across stages, while others, including LUSC, BRCA, LUAD/LUSC, and CESC, show no consistent trend. These findings indicate that the relationship between POLE2 expression and cancer stage is not universal but may reflect distinct biological roles of POLE2 in different tumor types.

### POLE2 promoter methylation analysis

One of the ways to investigate gene expression activity is to look at the methylation level of the gene promoter. We used HumanMethylation450 BeadChip data, which targets 96% of CpG islands in the human genome [88], that was part of the TCGA project. Combined methylation levels of all CpG sites (Fig. 6A) revealed that the POLE2 promoter is significantly hypermethylated (Δβ = 0.024) in cancer samples (n=8281) compared to normal samples (n=723). Upstream to POLE2, there are 12 CpG cites: cg11095277, cg10038907, cg09325174, cg10195098, cg26030540, cg15522118, cg15455399, cg23418097, cg23761011, cg04379738, cg27022570, and cg3496872 (Fig. 6B). If we analyze methylation level of each CpG site separately, we see that the predominant part of them are hypermethylated, while hypomethylations in only two sites are observed. Among the hypermethylated CpG sites, cg04379738 demonstrated the most substantial methylation difference between cancer and normal tissues (Δβ = 0.107), indicating meaningful hypermethylation in cancer. The differences at all other sites were below 0.1, and therefore are not considered biologically meaningful despite being statistically significant. (Fig. 6B). CpG sites cg11095277 (Δβ = –0.008) and cg3493872 (Δβ = –0.021) showed hypomethylation in cancer, but the differences were below 0.1 and therefore not biologically meaningful. Additionally, these two sites are not located within CpG island 52, based on data from the UCSC Genome Browser (Fig. 6C), which can mean that they are less associated with the POLE2 promoter region. Although the POLE2 promoter has not been definitively characterized, the “EPDnew v6” track [89] in the UCSC Genome Browser identifies an experimentally supported promoter, POLE2_1 (chr14:50154922–50154981, 60 bp in length). In contrast, the “Genome Segmentation by combined Segway+ChromHMM from the ENCODE/Analysis” track [90] defines broader regulatory regions, including a TSS region (Predicted promoter region including transcription start site) and a PF (Predicted promoter flanking region) region. These span from chr14:50152604 to chr14:50155774 across various cell lines, covering a 3,170 bp fragment. However, this extended region does not include the CpG sites cg11095277 and cg3493872; therefore, these sites were excluded from the promoter region in further analyses.

**Figure 6.**
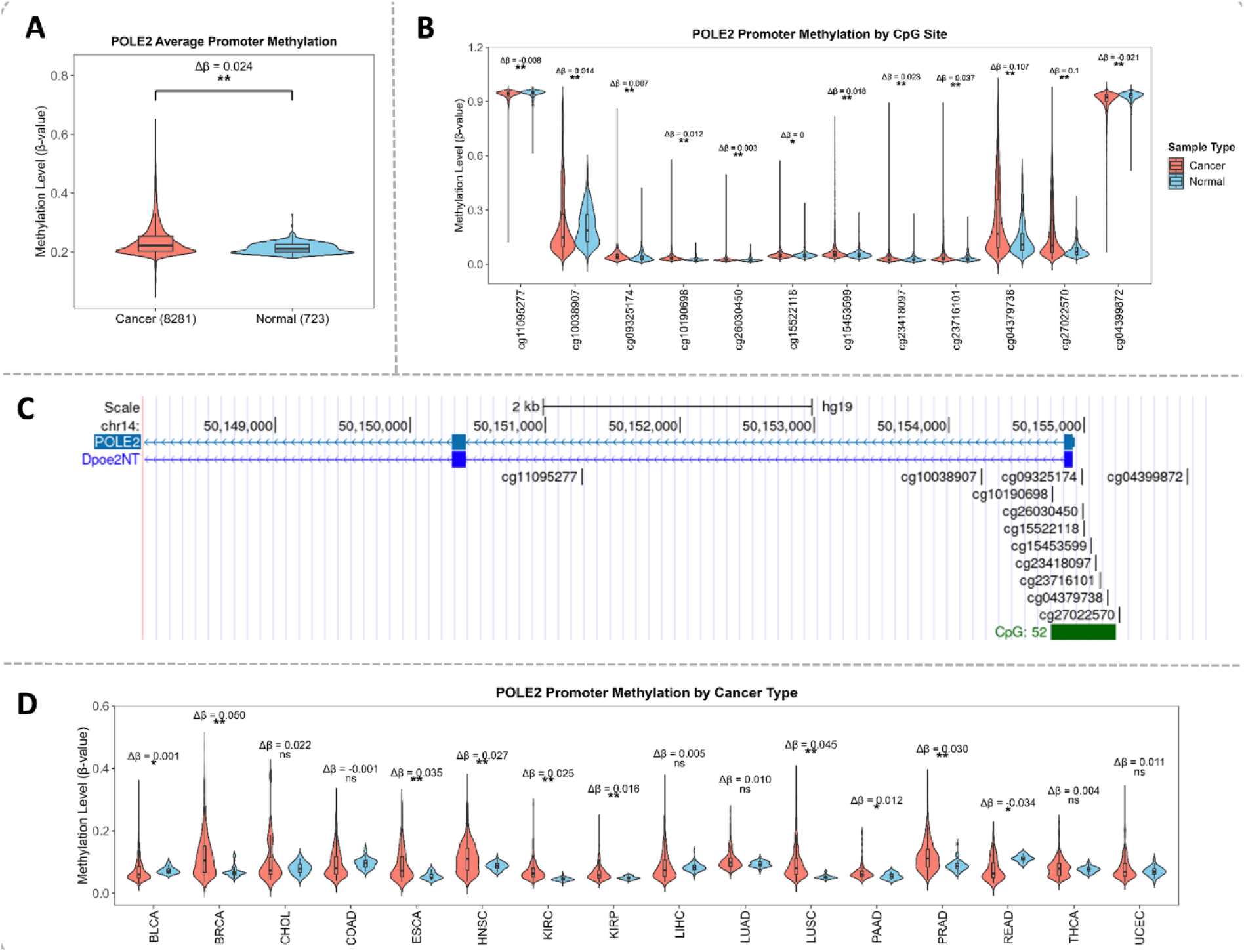
Comparison of POLE2 CpG site methylation levels in cancer and normal samples based on Illumina HumanMethylation450 BeadChip (450K array) data. **A.** Average POLE2 methylation levels across TCGA cancer samples (n = 8,281) and TCGA normal samples (n = 723). **B.** Methylation levels at individual CpG sites within the POLE2 gene region, shown separately for cancer and normal tissues. Numbers represent mean methylation difference (Δβ) -difference in average β-values between cancer and normal groups. **C.** UCSC Genome Browser view of the POLE2 locus, with CpG probe locations from the 450K methylation track. **D.** Average POLE2 promoter methylation levels for CpG sites within the POLE2 promoter region. Statistical significance was assessed using the Wilcoxon rank sum test (Mann–Whitney U test). Significance thresholds: p ≤ 0.01 (**), p ≤ 0.05 (*).

Next, POLE2 promoter methylation levels were assessed for 16 TCGA cancer types (Fig.6D). Statistically significant increases in methylation were observed in BRCA (Δβ = 0.050), ESCA (Δβ = 0.035), HNSC (Δβ = 0.027), KIRC (Δβ = 0.025), KIRP (Δβ = 0.016), LUSC (Δβ = 0.045), PAAD (Δβ = 0.012), PRAD (Δβ = 0.030), BLCA (Δβ = 0.001), and READ (Δβ = –0.034). However, the extent of these differences remained modest across all cases and is unlikely to reflect a substantial biological impact. In contrast, CHOL (Δβ = 0.022), COAD (Δβ = –0.001), LIHC (Δβ = 0.005), LUAD (Δβ = 0.010), THCA (Δβ = 0.004), and UCEC (Δβ = 0.011) showed no statistically significant differences in methylation between cancer and normal samples. Given that the overall differences in promoter methylation are minimal—even among statistically significant cases—it is unlikely that epigenetic regulation through DNA methylation is a major contributor to the upregulation of POLE2 mRNA expression in cancer. Nevertheless, the promoter region is predominantly hypermethylated in cancer samples. To further investigate the potential relationship between methylation and gene expression, we performed Pearson correlation analysis between POLE2 promoter methylation and mRNA expression levels (Fig. S4). This analysis revealed a weak but statistically significant negative correlation in cancer samples (r = –0.20), indicating that higher methylation is modestly associated with lower POLE2 expression. In contrast, the correlation in normal samples was very weak (r = –0.06) and only marginally significant, suggesting that DNA methylation likely plays a limited regulatory role in controlling POLE2 expression, particularly in the cancer context.

The correlation between methylation and expression was calculated separately for each cancer to investigate whether this effect is consistent across cancer types. Several tumor types showed notable correlations. TGCT exhibited the strongest negative correlation (r = –0.49), followed by LIHC (r = –0.24) and CESC (r = –0.13), indicating that increased methylation is associated with reduced POLE2 expression in these cancers. In contrast, LUSC (r = 0.12) and ESCA (r = 0.16) showed positive correlations, suggesting that the relationship may vary across cancer types and is not strictly repressive.

In normal adjacent tissues, fewer cancer types showed significant correlations, likely due to smaller sample sizes. Still, positive correlations were observed in BRCA (r = 0.24), LIHC (r = 0.55), and BLCA (r = 0.63).

In summary, the mean difference in promoter methylation between cancer and normal tissues is generally slight. However, the consistent hypermethylation observed across most CpG sites and in average comparisons suggests the involvement of additional regulatory mechanisms in POLE2 promoter control or potential inaccuracies in promoter region annotation. A positive correlation between promoter methylation and mRNA expression in normal tissues may reflect non-canonical regulation, such as activity from non-CGI or alternative promoters where methylation does not repress transcription. A weaker positive correlation in cancer may indicate disrupted epigenetic control or partial loss of regulatory stability [91].

### Isoform expression analysis

Alternative splicing (AS) generates multiple mRNA isoforms from a single gene, which are often regulated in a tissue- and cancer-type-specific manner. Many cancer-associated isoforms arise due to aberrant splicing, some of which are unique to tumors. Because of this specificity, isoform-level expression can serve as a biomarker for cancer diagnosis, prognosis, and treatment selection more precise than total gene expression [92]. Here, we studied transcript’s abundance relative to the total POLE2 expression (Fig. 7A, Fig. S5) and their structure (Fig.7B). Notably, the non-coding isoform POLE2-010 (7 exons) emerges as the most abundant isoform, representing approximately 32% of total POLE2 abundance in cancer samples, and similarly high proportions in Normal_GTEX (36%) and Normal_TCGA tissues (29%). This high abundance of a non-coding isoform is particularly noteworthy, suggesting a significant, potentially regulatory role despite lacking protein-coding capacity. Among coding isoforms, POLE2-001 (19 exons) is highly represented, comprising around 28% of total POLE2 abundance in cancer tissues, slightly increasing to 31% in Normal_GTEX and 35% in Normal_TCGA. Another coding isoform, POLE2-012 (17 exons), also shows significant abundance, consistently around 25% across cancer and normal samples, highlighting stability in its relative expression. Moderately abundant coding isoforms include POLE2-008 (3 exons), with reduced representation in cancer (11%) compared to Normal_GTEX (17%) and Normal_TCGA (14%), and POLE2-002 (18 exons), which notably decreases from around 13-14% in normal tissues to approximately 6% in cancer. Lower-abundance of non-coding isoforms POLE2-003, POLE2-006, and POLE2-007 (3 exons) demonstrate clear reductions in cancer samples compared to normal tissues. Similarly, problematic isoforms POLE2-004 (2 exons), POLE2-005 (3 exons), and POLE2-011 (4 exons), characterized by lower expression and possible structural or annotation issues, consistently show reduced abundance in cancer tissues relative to normal tissues. The analysis of POLE2 isoform percentages indicates that the previously observed overexpression of the POLE2 gene in cancer primarily results from increased abundance of two coding isoforms (POLE2-001 and POLE2-012) and one prominent non-coding isoform (POLE2-010). Given that total isoform percentages sum to 100%, the decreased relative proportions of problematic and other less abundant isoforms in cancer tissues likely reflect their dilution due to the substantial upregulation of these dominant isoforms, rather than an actual decrease in their absolute expression levels. This highlights that isoform-specific regulation contributes significantly to the elevated expression of POLE2 observed in cancer.

**Figure 7.**
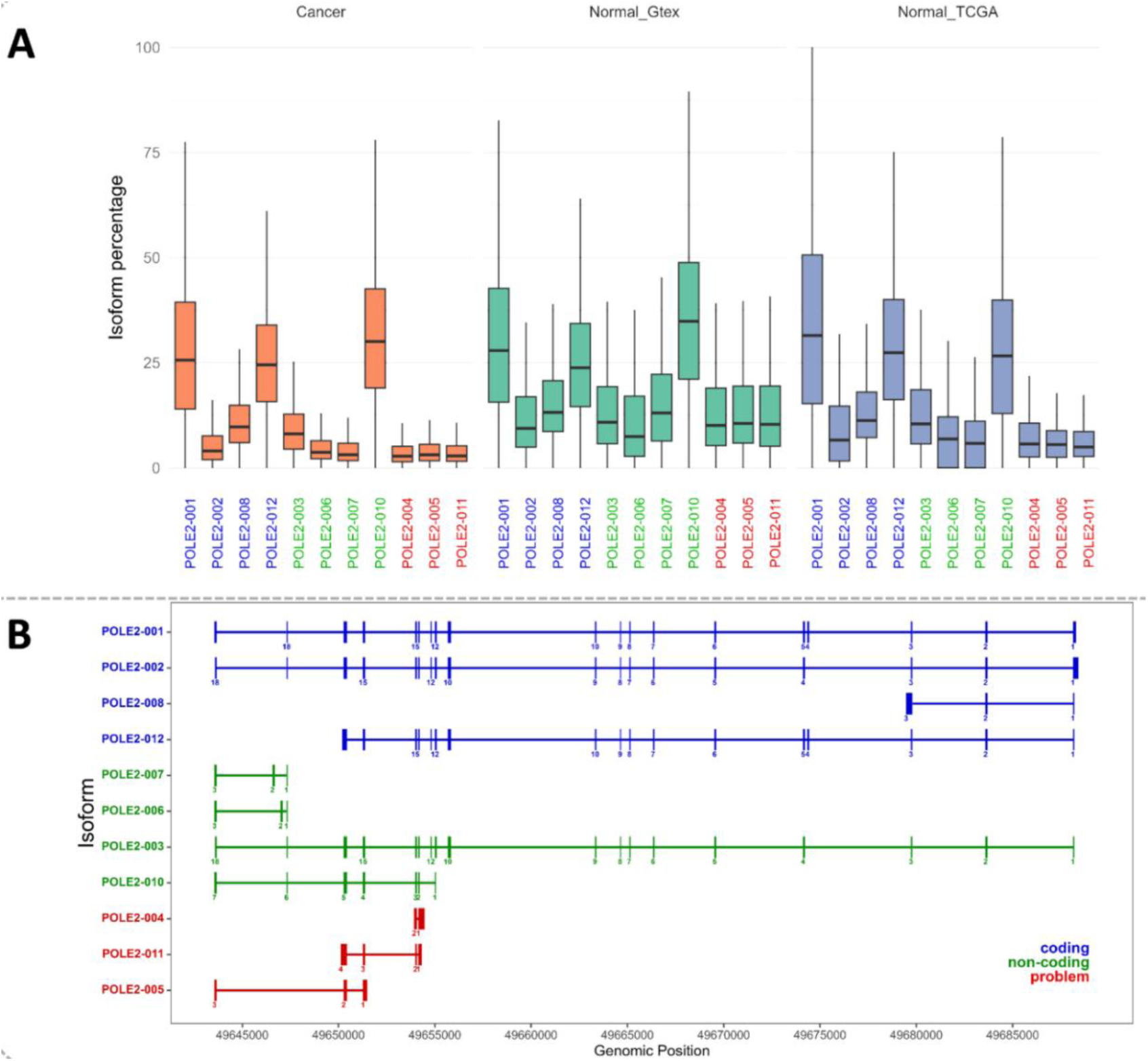
POLE2 isoform expression and transcript structure in cancer and normal tissues. **A.** I**soform percentage (IsoPct)** of each POLE2 transcript across three sample types: cancer(n = 9,163), GTEx normal(n = 5,354), and TCGA normal(n = 712), derived from TOIL-processed isoform expression data obtained via the UCSC Xena database. IsoPct represents the proportion of a transcript’s abundance relative to the total expression of its parent gene (POLE2), allowing us to assess **isoform usage** rather than absolute expression levels. **B.** Exon–intron structures of 11 POLE2 transcript isoforms aligned to the same genomic locus, based on UCSC Genome Browser annotations. Each horizontal line represents a transcript; vertical blocks indicate exons, and connecting lines represent introns. Isoforms are color-coded by classification: coding (blue), non-coding (green), and problematic/incomplete annotations (red). Significance thresholds: p ≤ 0.01 (**), p ≤ 0.05 (*).

Focusing on cancer-specific isoforms can enhance diagnostic precision, and we have found that coding POLE2-001 and POLE2-012, as well as non-coding POLE2-010, are predominantly expressed in cancer. Additionally, isoform-specific expression can explain why promoter methylation and gene expression are not inversely correlated. For example, non-coding isoform POLE2-010, whose exons are located distant from the hypermethylated POLE2 promoter region, has a bigger proportion in isoform abundance, and can partially explain the POLE2 expression rise during promoter hypermethylation. However, further studies on isoform-specific promoter methylation are needed to draw definitive conclusions.

### POLE2 protein levels in cancer samples do not always correspond to their mRNA expression levels

The comparison between mRNA and protein expression levels across seven cancer types—BRCA, GBM, LIHC, LUAD, LUSC, PAAD, and UCEC—was conducted using data from the UALCAN database (Fig. 8). These cancer types were selected because CPTAC mass spectrometry data with profiled POLE2 protein level was available only for a limited number of cancer types.

**Figure 8.**
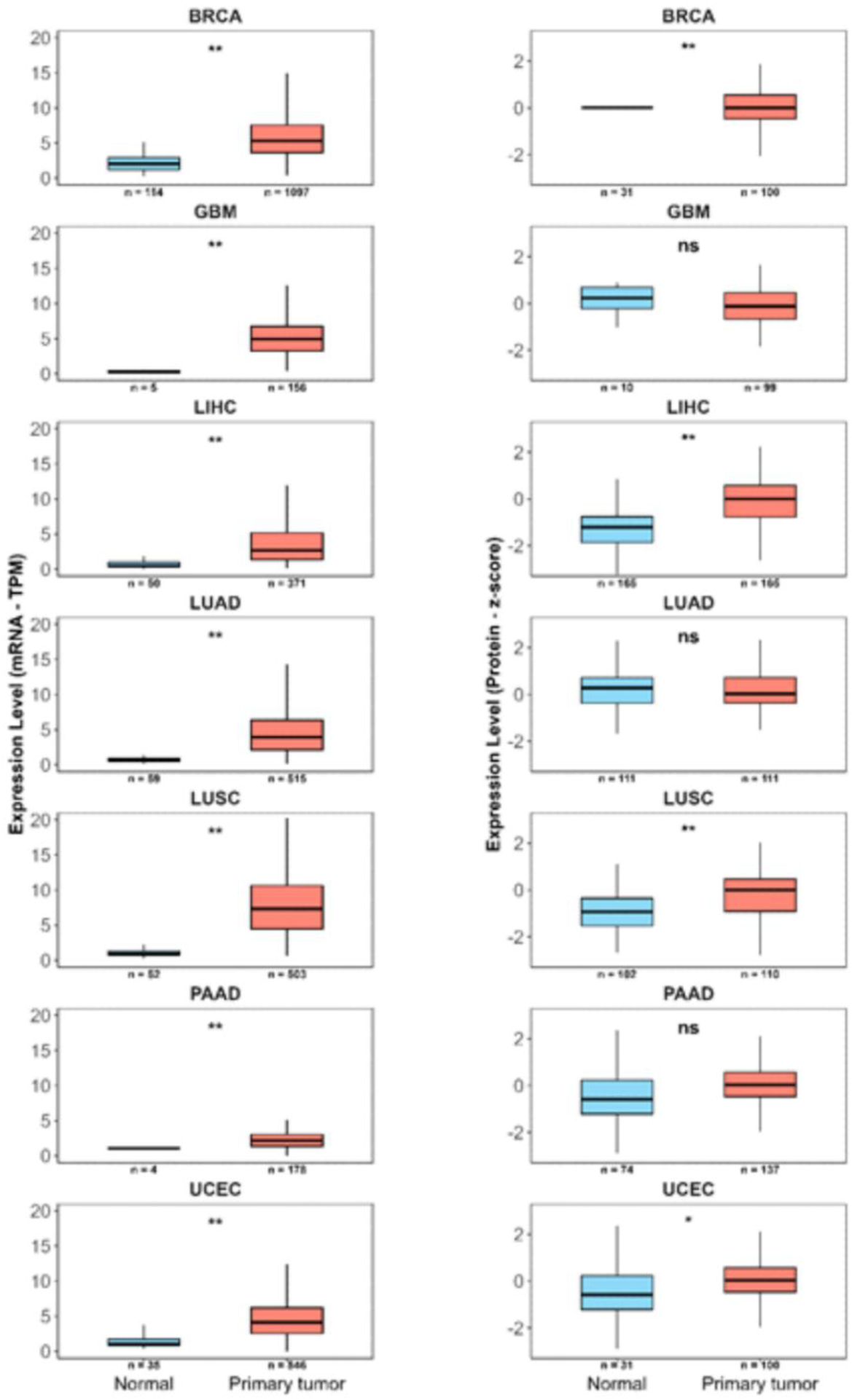
Comparison of POLE2 mRNA and protein expression in cancer versus normal tissues based on the UALCAN database. Boxplots on the left show POLE2 mRNA expression levels (TPM) across normal and primary tumor tissues in selected cancer types. Boxplots on the right represent corresponding POLE2 protein expression levels (z-scores) in the same tissue types. Statistical significance was calculated using the Wilcoxon rank sum test (Mann–Whitney U test). Significance levels: *p* ≤ 0.01 (**), *p* ≤ 0.05 (*).

In BRCA, mRNA levels in tumors (n=1097, median = 5.32) were significantly higher than in normal tissues (n=114, median = 2.02), and this difference was also seen at the protein level (tumor n=100, median = 0.24; normal n=31, median = -0.19). GBM showed a strong increase in mRNA (tumor n=156, median = 4.92; normal n=5, median = 0.26), but no significant change in protein (tumor n=99, median = 0.02; normal n=10, median = -0.06). In LIHC, transcript levels rose notably (tumor n=371, median = 5.31; normal n=50, median = 0.35), and protein values also increased (tumor n=99, median = 0.53; normal n=10, median = -0.36). LUAD showed elevated mRNA levels in tumors (n=515, median = 4.97) compared to normal (n=59, median = 0.32), but protein levels remained similar (both groups n=165, medians = 0.08 and 0.03), with no significant difference. In LUSC, mRNA was strongly upregulated (tumor n=503, median = 7.27; normal n=52, median = 0.28), and protein levels also increased (tumor and normal n=111, medians = 0.38 and -0.23). PAAD displayed a rise in transcript levels (tumor n=178, median = 1.05; normal n=4, median = 1.07), though the difference was small, and protein values showed no significant variation (tumor n=137, median = 0.12; normal n=74, median = -0.10). In UCEC, transcript levels increased clearly (tumor n=546, median = 4.38; normal n=35, median = 0.56), and protein also showed a notable shift (tumor n=100, median = 0.27; normal n=31, median = -0.19).

Across all examined cancers, the magnitude of change in mRNA levels between tumor and normal tissues was consistently greater than that observed at the protein level. BRCA, LIHC, LUSC, and UCEC showed significant changes in both data types. In contrast, GBM, LUAD, and PAAD displayed increased mRNA without significant differences in protein levels. It can be partially explained by the higher abundance of non-coding POLE2-010 isoform in cancer, as described above, as well as translational efficiency, protein stability, and sensitivity of the mass spectrometry method utilized in that study.

### Mutations in POLE2 identified in cancer samples

POLE2 encodes a core structural subunit of the DNA polymerase epsilon complex, which is essential for high-fidelity leading-strand DNA synthesis during DNA replication. Although POLE2 lacks enzymatic activity, mutations in this gene can compromise replication accuracy by destabilizing the polymerase complex or impairing its interaction with DNA. Analysis of somatic mutation data from the COSMIC database [93] and previous literature has revealed recurrent POLE2 alterations across several cancer types, including colorectal, endometrial, renal, and biliary tract tumors. These mutations include nonsense variants, frameshift mutations, and missense changes, some of which are predicted to disrupt protein structure or hinder complex formation. Table S5 and the lollipop plot in Fig. 9 summarize all POLE2 mutations, with a recurrence count of >1, curated from the COSMIC database, highlighting their distribution and functional classification across the gene.

**Figure 9.**
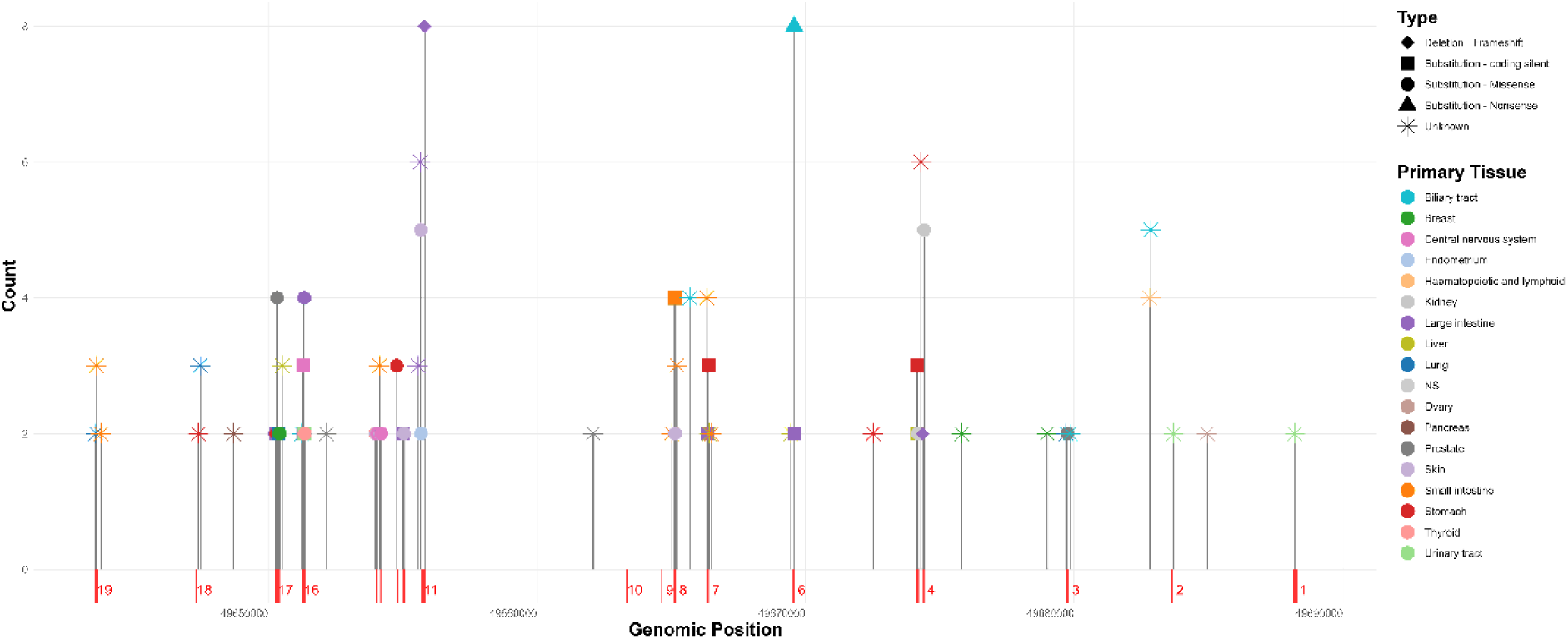
Distribution of POLE2 mutations across exons and introns. Lollipop height indicates mutation frequency in cancer samples. Colors represent tissue of origin, and shapes denote mutation types. Exons are shown as red blocks; introns span between them.

To assess the spatial distribution of POLE2 mutations, the protein structure was visualized using the cryo-EM structure available in the Protein Data Bank (PDB ID: 7PFO). COSMIC-annotated mutations were mapped onto this structure [94] using ChimeraX software. Mutations with a recurrence count greater than four were highlighted to identify regions of potential functional relevance (Fig. S6). Notably, a clustering of mutations was observed in a specific region of the protein, suggesting a potential hotspot for structural or functional disruption. This region was marked by a higher local density of missense and truncating mutations, which may interfere with POLE2’s interaction within the polymerase epsilon complex. The structural visualization provides additional context for understanding how specific alterations may contribute to replication dysfunction and cancer development.

A germline POLE2 variant c.823C>A was reported in five individuals from the same family affected by colorectal cancer, along with a frameshift mutation, c.1328_1329insT (p.Leu443Phefs*17), predicted to disrupt protein function [95]. Subsequently, another frameshift mutation, c.1406dupT (p.Leu469PhefsTer17) was identified in four samples from colorectal cancer patients [96]. Later, additional germline POLE2 variants, including c.655A>G (one heterozygous carrier), c.1414T>C (two heterozygous carriers), c.50G>C (one carrier), and c.856A>G (one carrier), were reported [97]. Although many of these alterations are currently classified as variants of uncertain significance (VUS), their recurrence in colorectal cancer cohorts and their potential to impair replication fidelity suggest a possible role in promoting genomic instability and tumor development. Germline mutations found in these studies are summarized in Table 1.

**Table 1.** Germline mutations identified in POLE2.

| Protein change | Mutation type | Nucleotide change | Tissue and cancer type | Reference |
| --- | --- | --- | --- | --- |
| p.R17P | Substitution - Missense | c.50G>C | Colorectal carcinoma | [97] |
| p.T219A | Substitution - Missense | c.655A>G | Colorectal carcinoma | [97] |
| p.M286V | Substitution - Missense | c.856A>G | Colorectal carcinoma | [97] |
| p.L275I | Substitution - Missense | c.823C>A | Colorectal carcinoma | [95] |
| p.Y472H | Substitution - Missense | c.1414T>C | Colorectal carcinoma | [97] |
| p.L443Ffs*17 | Frameshift- Insertion | c.1328_1329insT | Colorectal carcinoma | [95] |
| p.L469FfsTer17 | Frameshift- duplication | c.1406dupT | Colorectal carcinoma | [96] |

### Enrichment Analysis of POLE2-Correlated Genes in Cancer

In order to investigate molecular mechanisms of POLE2 overexpression and association with cancer, we studied genes that are highly correlated (Pearson correlation value- PCC >0.8) with POLE2 and therefore have similar expression patterns. To obtain convincing results, we used four databases as a source of POLE2 correlated genes and overlapped them using an UpSet plot.

The UpSet plot (Fig. 10) illustrates how genes are distributed across five datasets: GTEx_Normal and Analyzer_Normal for normal tissue, and GEPIA_Cancer, Analyzer_Cancer, and UALCAN_Cancer for tumor tissue (Fig. 10, Fig. S7, Fig. S8). The total number of gene occurrences across all sets is 2303 (Set Size Total), while 1656 genes fall into exclusive intersections (Intersection Size Total), indicating the number of unique genes assigned to specific dataset combinations.

**Figure 10.**
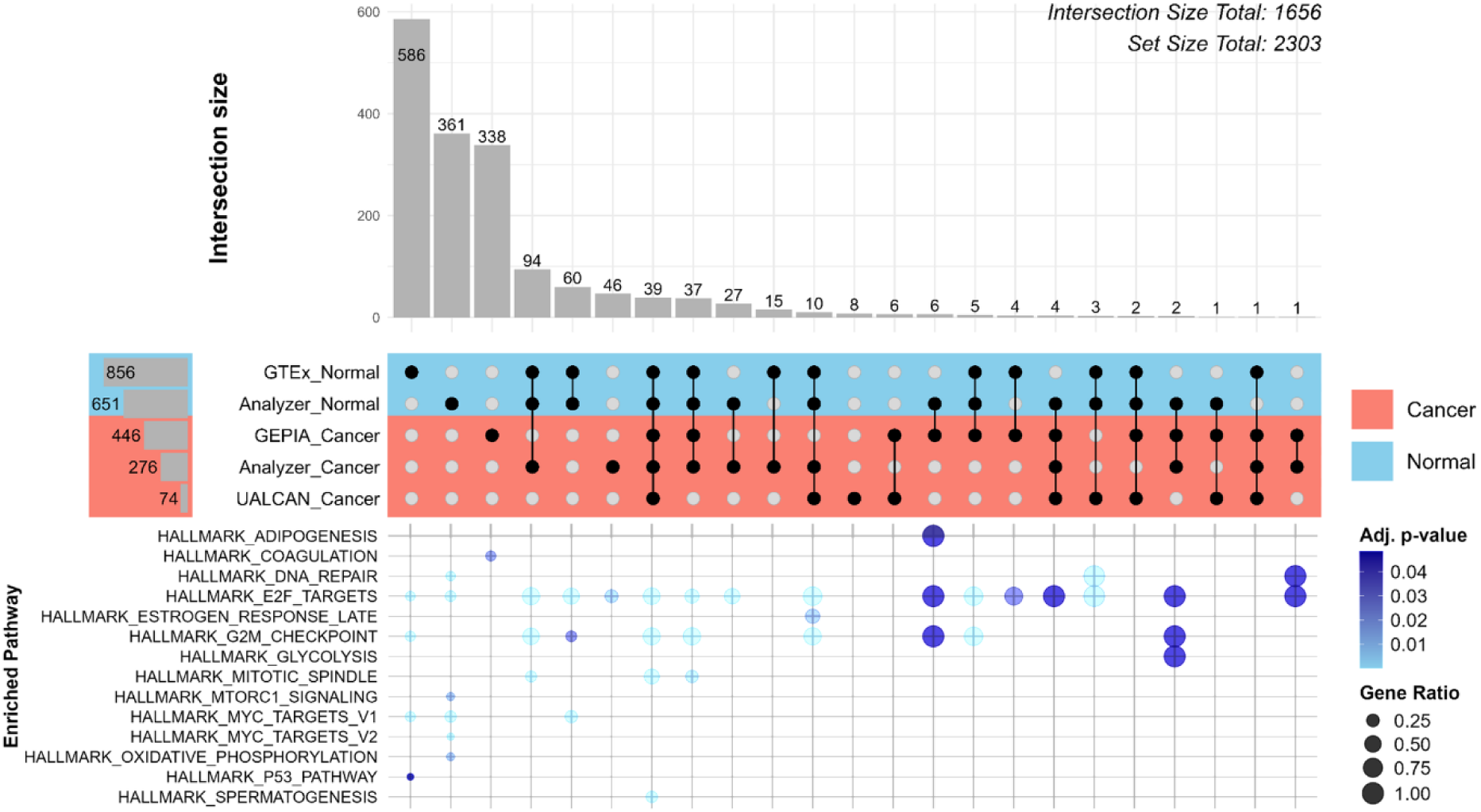
Overlap of genes highly correlated with POLE2 across multiple datasets in cancer and normal tissues. UpSet plot showing the intersection of genes with a Pearson correlation coefficient > 0.8 with POLE2 expression, derived from six datasets from 4 independent databases: GTEx, GEPIA, UALCAN, and AnalyzeR, each separated by sample type. Top bars indicate the number of overlapping genes shared among dataset combinations, while the left bars show the total number of highly correlated genes per dataset. “Set Size Total” represents the total number of gene occurrences across all input sets, counting duplicates when a gene appears in multiple sets. In contrast, “Intersection Size Total” reflects the number of unique genes that are assigned to exclusive intersections—each gene counted only once based on its specific combination of set membership. Dot plot representing the results of Over-Representation Analysis (ORA) performed on intersecting POLE2-correlated genes using the **HALLMARK gene set collection.** Dot size reflects the gene ratio, and color intensity indicates statistical significance (adjusted *p*-value).

Cancer-specific exclusive intersections are led by “GEPIA_Cancer”, which alone contains 338 unique genes, making it the largest single-source cancer gene set. Less prevalent cancer-only intersections include “Analyzer_Cancer” with 46 genes and the combined “UALCAN_Cancer. GEPIA_Cancer” with six genes. Functional enrichment analysis of these cancer-specific groups reveals that “GEPIA_Cancer” genes are enriched in the hallmark coagulation pathway (adjusted p = 0.024), suggesting potential links to tumor-associated clotting mechanisms. “Analyzer_Cancer” genes are enriched in E2F targets (p = 0.013), indicating involvement in cell cycle regulation.

For normal-specific intersections, the largest group is “GTEx_Normal” with 586 unique genes, followed by “Analyzer_Normal” with 361 genes, and their shared intersection “Analyzer_Normal. GTEx_Normal” containing 60 genes. These normal cell-related groups are strongly enriched in proliferation-associated pathways. “GTEx_Normal” genes are enriched in G2M-checkpoint, E2F, and MYC targets, indicating active regulation of mitosis and transcription. “Analyzer_Normal” also shows strong enrichment for MYC and E2F targets, further supporting that the normal cell-specific expression includes tightly regulated growth-associated programs.

Mixed intersections—those involving combinations of both cancer and normal datasets—contribute smaller gene groups compared to the large single-source intersections, but show a high density of functional enrichment. For instance, the intersection “Analyzer_Cancer. Analyzer_Normal. GTEx_Normal” contains 94 genes and is enriched in multiple hallmark pathways, including E2F targets and G2M checkpoint, reflecting strong involvement in cell cycle regulation. Similar enrichment is observed in the five-way intersection that includes all datasets and contains 39 genes, and the two four-way intersections containing 37 and 10 genes, respectively (Fig. 10).

From the comparison of the enrichment results across cancer-specific, normal-specific, and mixed intersections, distinct patterns emerge. Normal cell-specific gene sets are strongly enriched in core proliferative and transcriptional programs, including E2F, G2/M checkpoint, and MYC targets. These reflect tightly regulated growth-associated functions active in normal tissues. Cancer cell-specific intersections are enriched in pathways such as coagulation and E2F targets, suggesting a shift toward deregulated cell cycle control and tumor-associated processes. Interestingly, mixed intersections involving both cancer and normal datasets also show strong enrichment in proliferation-related pathways, including E2F and MYC targets, and G2/M checkpoint. The enrichment of these pathways in intersections containing both normal and cancer datasets may suggest that such genes are part of essential regulatory networks that remain active across states or are particularly responsive to changes in POLE2 expression. Their consistent correlation with POLE2 in cancer implies that when POLE2 is dysregulated, these genes may be co-altered as part of broader transcriptional shifts.

To identify genes strongly correlated with POLE2 in a cancer-specific manner and to explore patterns that may distinguish one TCGA cancer type from others, an intersection analysis was performed across all available TCGA cancer types using five expression sources: GEPIA_Cancer, UALCAN_Cancer, Analyzer_Cancer, GTEx_Normal, and Analyzer_Normal. Since Correlation AnalyzeR and GTEx datasets are tissue-based and not cancer-type-specific, their mappings to TCGA cancer types were adapted using the cancer classification used in GEPIA2, with minor adjustments to ensure consistency.

Initially, 23 TCGA cancer types were available across the datasets. However, to ensure meaningful intersection analysis, only cancer types with at least three represented datasets were included. As shown in the Fig. S8, the final selection includes ten cancer types: KICH, KIRC, KIRP, LAML, STAD, SKCM, LUAD, ESCA, DLBC, and GBM.

The genes found in cancer-specific intersections (Fig. S8) were subject to enrichment analysis independently for each cancer type to explore their associated biological functions (Fig 11).

**Figure 11.**
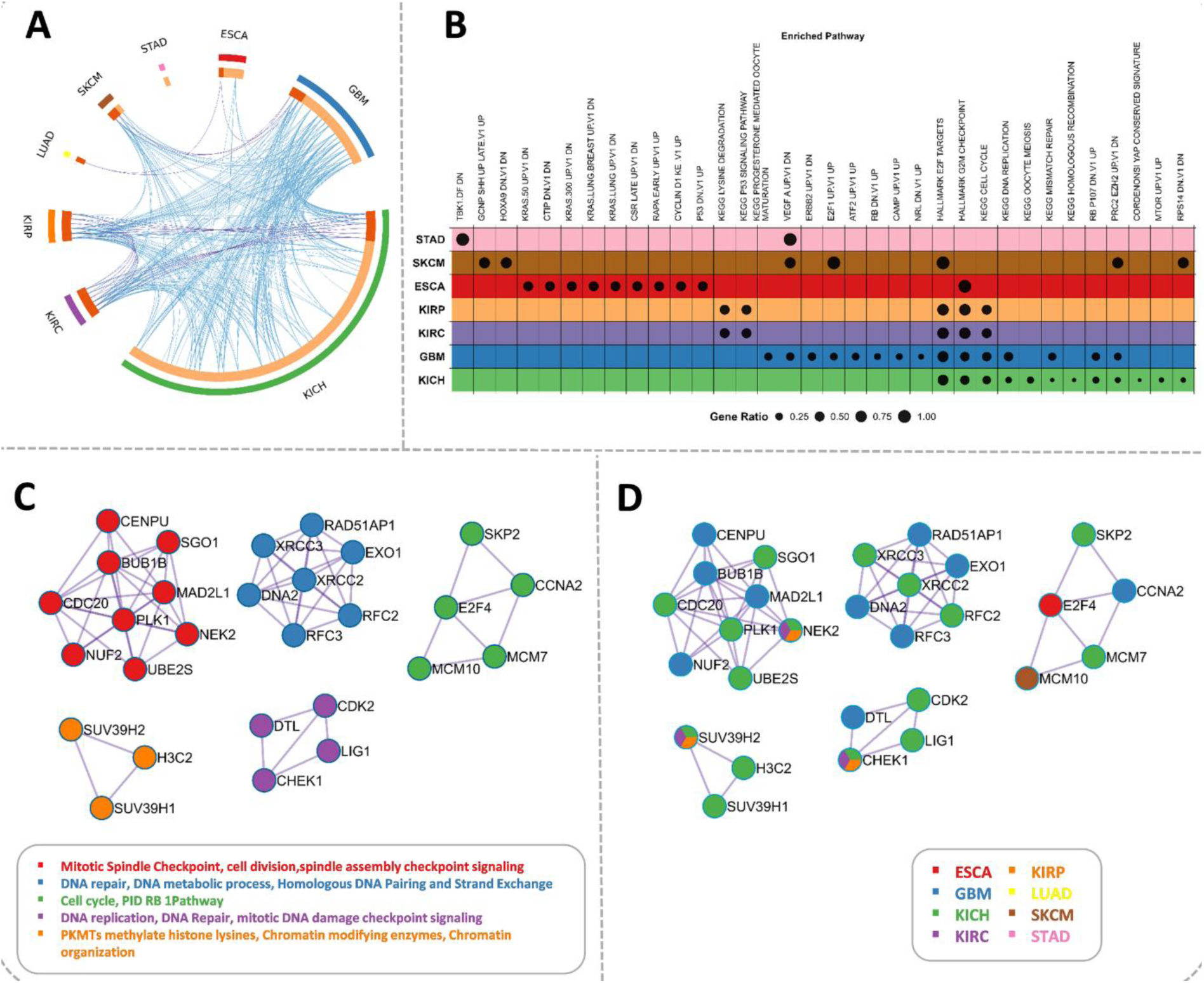
Enrichment analysis of gene groups exclusively correlated with POLE2 in cancer. A-. Circos plot retrieved from Metascape showing overlaps among genes that share the same enriched ontology term(s). Purple curves connect identical genes, while blue curves link genes associated with the same ontology term. **B-**Overexpression analysis results of gene groups exclusively correlated with POLE2 in cancer. Only significant categories (p-value<0.05) are shown. Gene Ratio refers to the proportion of input genes associated with a particular enriched term. **C-** Protein-protein interaction (PPI) network from Metascape for all cancer types merged and colored by MCODE(Molecular Complex Detection) functional clusters. **D**- PPI network from Metascape for all cancer types merged and colored by cancer types – names of initial gene groups

The Circos plot (Fig. 11A) illustrates shared genes among the POLE2-correlated gene sets from different cancer types. Many shared genes are observed among the three kidney cancers—KIRC, KIRP, and KICH—which reflects their common tissue origin, as they were all mapped from kidney-specific expression data. GBM and KICH appear to share the highest number of POLE2-corralated genes with other cancer types.

The dot plot (Fig. 11B) summarizes the over-representation analysis (ORA) results for gene sets exclusively correlated with POLE2. While enrichment analysis was performed for 11 cancer types, only 7 of them are represented in the plot, as they showed significant enrichment in at least one pathway. The enrichment analysis reveals distinct biological processes across several cancer types. Individual enriched genes from overexpression analysis for each cancer type are shown on the sunburst plot (Fig. S9).

Gene sets related to DNA replication and repair include DNA replication and mismatch repair for GBM and KICH, as well as homologous recombination processes for KICH. However, each cancer type has different sets of enriched genes in the same term. For example, DNA replication in GBM contains PRIM1, DNA2, and RFC3, while KICH is enriched with LIG1, RNASEH2A, MCM7, and RFC2. As for mismatch repair, in GBM it includes RFC3 and EXO1, while in KICH it includes RFC2 and LIG1. Homologous recombination in KICH includes XRCC2 and XRCC3.

Enrichment of the gene set linked to epigenetic regulation (“PRC2 EZH2 UP.V1 DN”) [98] was observed in GBM, KICH, and SKCM. This set consists of the 200 most downregulated genes following EZH2 knockdown. In GBM, the overlapping genes include PRIM1, DLGAP5, CENPU, and DNA2. In KICH, the enriched genes are OIP5, SKP2, XRCC3, APLP1, and PSRC1, while in SKCM, the overlapping genes are MCM10 and DLGAP5.

PRC2 (Polycomb Repressive Complex 2) is a key epigenetic regulator involved in transcriptional repression through histone modification. EZH2 is the catalytic subunit of PRC2, responsible for histone methylation at lysine 27 (K27) on histone H3 and lysine 26 (K26) on histone H1 [98]. In GBM, KICH, and SKCM, gene sets that are downregulated upon EZH2 knockdown were found to be enriched. These genes are highly correlated with POLE2 expression and are positively regulated by EZH2, suggesting a potential coordination between POLE2-associated transcriptional programs and EZH2-mediated chromatin regulation in these cancers.

Another gene set linked to epigenetic regulation and associated with ribosomal function and stress response (“RPS14 DN.V1 DN”) [99] was enriched in KICH and SKCM. This set includes genes downregulated upon RPS14 knockdown. In KICH, the overlapping genes are SUV39H2, ESPL1, CDC20, and PLK1. In SKCM, the enriched genes include MCM10 and DLGAP5. SUV39H2 was also identified as a correlated gene of the lysine degradation pathway enriched in both KIRC and KIRP. SUV39H2 encodes a histone methyltransferase involved in lysine modification and heterochromatin formation, linking the pathway to epigenetic regulation.

RPS14 (Ribosomal Protein S14) is a component of the 40S small ribosomal subunit and plays essential roles in ribosome biogenesis, protein synthesis, and broader cellular regulatory pathways. Its loss disrupts the formation of the small ribosomal subunit, causing nucleolar stress and activating the p53 pathway, which can lead to cell cycle arrest and apoptosis [100]. The enrichment of downregulated genes upon RPS14 knockdown was detected in KICH and SKCM. This could indicate that in those cancers, the POLE2-correlated transcriptional program includes genes that are normally suppressed under ribosomal stress, potentially reflecting a state of sustained proliferation despite p53 signaling or ribosomal disruption. In this context, POLE2 may be associated with transcriptional programs that help override growth-inhibitory responses, maintaining expression of mitotic regulators such as CDC20, PLK1, or MCM10 even under conditions that would typically lead to their downregulation.

The most abundant gene sets were associated with cell cycle regulation and mitotic control and enriched across different cancer types. GBM, KICH, KIRC, and KIRP show enrichment for E2F-targets, G2M checkpoint, and cell cycle processes. SKCM also show enrichment in E2F targets. In GBM and KICH, the “RB_P107_ DN.V1_UP” signature was also enriched. The p53 signalling pathway gene set was enriched in KIRC and KIRP, each with CHEK1 as a correlated gene.

Given that genes that exclusively correlate with POLE2 in cancer overlap with those upregulated after RB knockout, it can mean that those genes are normally inhibited by RB presence. This data confirms that the expression of POLE2 correlated genes is regulated by the E2F transcription factor.

ESCA shows enrichment for the G2M checkpoint gene set as well as the “CYCLIN_D1_KE.V1_UP” gene set, composed of genes upregulated in cells overexpressing a mutant form of Cyclin D1, and the “P53_DN.V1_UP” gene set, which includes genes upregulated in the context of TP53 loss or mutation. To summarize, this analysis highlights the involvement of POLE2-correlated genes in key cellular processes. Using cancer signature gene sets proved especially valuable, as it allowed observation of how these genes behave across different cancer types and experimental conditions. While many enrichment terms appeared significant in multiple cancers, the specific genes contributing to these terms varied considerably. This suggests that although POLE2 is associated with similar biological processes, the exact gene networks involved are cancer-specific.

As a part of functional enrichment in Metascape, the PPI network was constructed, which revealed five distinct clusters (Fig. 10C-D), each corresponding to core biological processes implicated in cancer.

The first cluster, enriched in mitotic spindle checkpoint and cell division genes, formed a tightly connected module. These genes are predominantly associated with kidney tumors, mostly KICH (CDC20, SGO1,PLK1,UBE2S,NEK2), as well as GBM (CENPU, BUB1B,MAD2L1,NUF2). CDC20 is functionally connected to all other genes in this module, suggesting that it might be the main functional driver for deregulation in cell division.

The second cluster involved genes regulating DNA repair, DNA metabolic processes, and homologous recombination. These were strongly linked to GBM (RAD51AP1,EXO1,DNA2,RFC3) and KICH (XRCC3,XRCC2,RFC2).

The third cluster, centered around cell cycle control and E2F regulation, showed overlap across cancers including GBM (CCNA2), KICH (SKP2, MCM7), ESCA (E2F4), and SKCM (MCM10).

The fourth module was enriched in DNA replication, DNA repair, and checkpoint signaling, particularly active in KICH (CDK2, LIG1, CHEK1), KIRC, KIRP (CHEK1), and GBM (DRL), reflecting their reliance on replicative stress responses. These genes partially bridge the DNA repair and cell cycle modules, reinforcing their role as key integrators of cell fate decisions.

Lastly, a distinct chromatin regulation cluster (e.g., SUV39H1, SUV39H2, H3C2) was specifically enriched in KICH, KIRP, and GBM. The lack of direct connections to other clusters may indicate that chromatin state changes act as independent modulators of oncogenesis in these contexts.

PPI network was done on genes exclusively and highly correlated with POLE2 across multiple cancers to uncover functionally coherent clusters involved in key processes like cell division, DNA repair, and chromatin regulation. This helped reveal cancer-specific patterns of gene co-regulation, offering insights into potential mechanisms of POLE2 involvement in tumor biology. Further investigation is needed to determine whether changes in POLE2 could influence these processes by altering the expression of the identified genes. This could help clarify POLE2’s functional role in tumor progression and its potential as a therapeutic target.

## CONCLUSIONS

This study integrates multiple data sources to provide more robust insights into how the expression of POLE2, the non-catalytic subunit of DNA polymerase ε, relates to survival and tumor progression across individual cancer types (Fig. 12). We conclude that POLE2 is consistently upregulated across multiple cancer types. Moreover, it stands out as the most overexpressed subunit within the group of replicative polymerases. Its expression levels, rather than copy number variation (CNV), emerge as a more reliable prognostic marker. Stage-dependent expression of POLE2 further suggests a role in tumor progression in certain cancers, such as ACC and KIRC, while others show less pronounced or no consistent trend. Additionally, analysis of mutations identified in POLE2 in cancer samples suggests that they cluster in a specific region, which could affect its interaction with the catalytic subunit and complex formation. This study also demonstrates that POLQ stands out as another polymerase with consistent upregulation, linked to replication stress tolerance. In contrast, several translesion polymerases are broadly downregulated.

**Figure 12.**
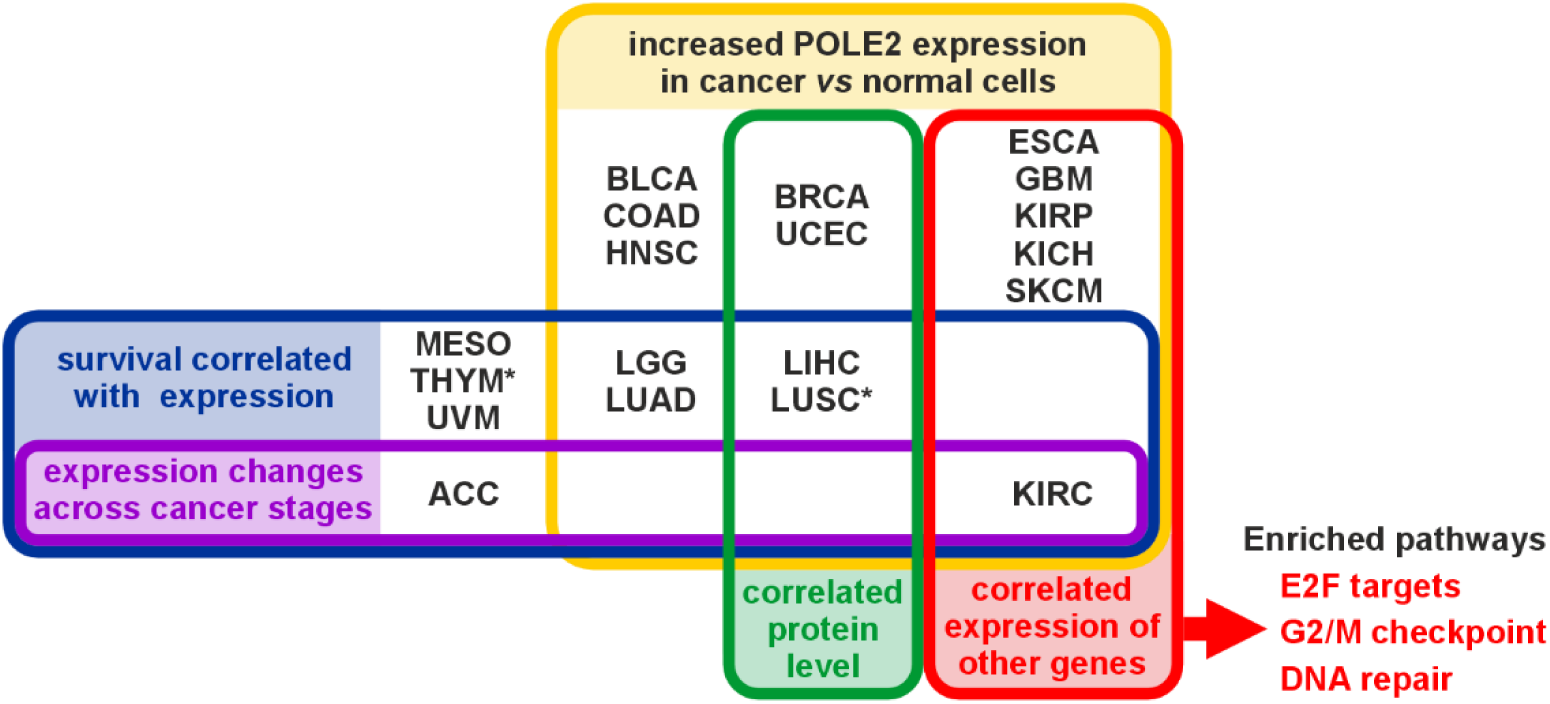
Summary of the most pronounced effects of POLE2 expression on cancer cells. POLE2 expression is upregulated across multiple cancer types (yellow). Moreover, depending on the cancer type, this correlates with a confirmed increase in protein level (green), a specific cancer stage (purple), or increased expression of other genes (red), mainly E2F targets and those involved in the G2/M checkpoint or DNA repair. In many cases, increased POLE2 expression correlates (blue) with poor survival prognosis, but in two cases [*] the effect is inverted.

Elevated POLE2 expression is associated with decreased overall survival in a cancer-type-specific manner, with particularly strong prognostic significance observed in ACC, LGG, LUAD, MESO, and potentially UVM, KIRC, and LIHC. In contrast, a protective effect was seen in THYM and LUSC, underscoring the context-dependent nature of its role. Despite CNVs influencing POLE2 expression, additional regulatory mechanisms, including promoter methylation, isoform usage, and epigenetic regulators like EZH2, contribute to its complex regulation. A consistent positive correlation between promoter methylation and gene expression, especially in normal tissues, suggests the presence of non-canonical regulation and highlights the need for isoform-specific epigenetic studies. Interestingly, POLE2 is tightly co-expressed with genes involved in DNA replication, cell cycle progression, and chromatin remodeling, and shows cancer-specific expression networks. Finally, network and enrichment analyses indicate that POLE2 is embedded in cancer-specific transcriptional programs that may help override normal growth-inhibitory signals, potentially mediated by E2F, EZH2, and RB pathways. These findings collectively position POLE2 as a functionally significant and clinically relevant marker, with a role that extends beyond DNA replication, and warrant further investigation into its biological function and therapeutic potential in oncology.

## AUTHOR CONTRIBUTIONS

Conceptualization: AK, DC, IJF, MD; Data Curation: AK, DC, IJF, MD; Formal Analysis: AK, DC, IJF, MD; Validation: AK, DC; Visualization: AK, DC; Writing – Original Draft Preparation: AK, DC, MD; Writing – Review & Editing: IJF, MD.

