## Supplementary material for "Pan-cancer multi-omics analysis identifies non-canonical oncogenic functions beyond DNA replication of the POLE2 subunit of DNA Polymerase epsilon": Figures S1, S2, S3, S4, S5, S6, S7, S8, S9

### SUPPLEMENTARY FIGURES

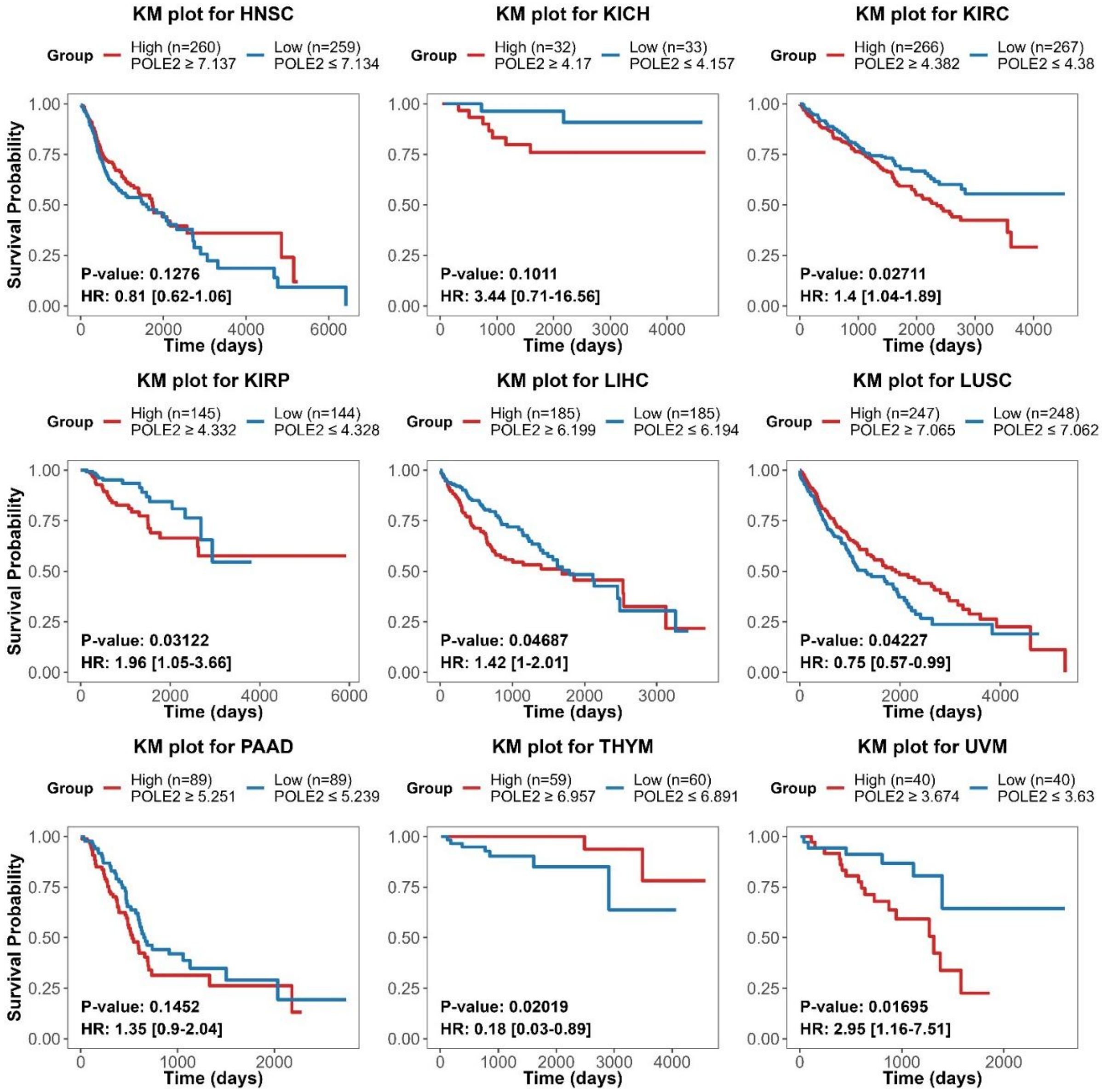


**Figure S1.** Kaplan–Meier plots from TCGA Pan-Cancer (PANCAN) data for selected cancer types from Fig. 1B, not shown in Fig. 1C. The plot displays the hazard ratio (HR) followed by its 95% confidence interval in square brackets.

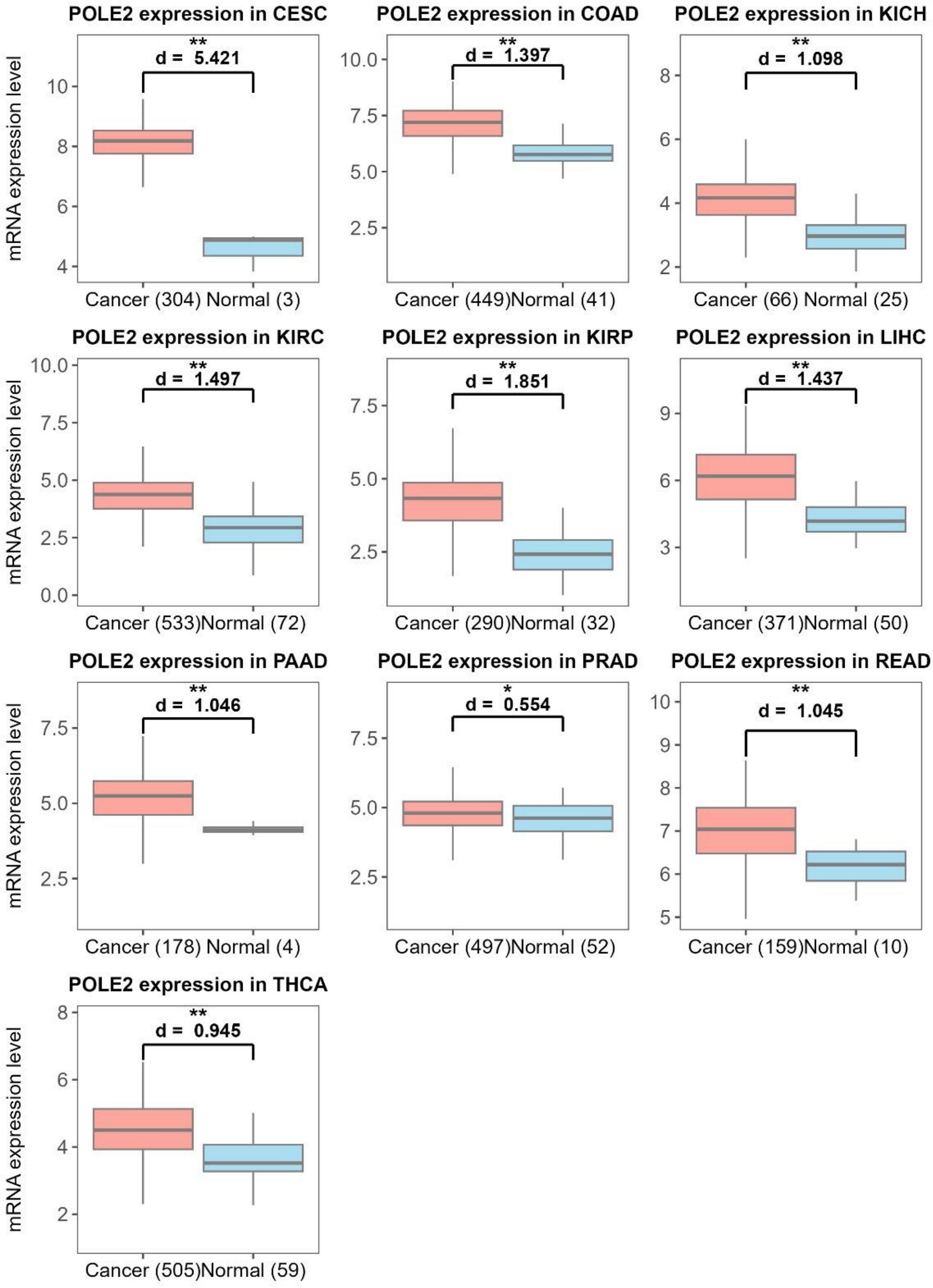


**Figure S2. .** Boxplots of POLE2 expression based on TCGA RNA-seq data for selected cancer types with consistent significance in at least three databases shown in Fig. 3A. Each plot displays the standardized mean difference (Cohen's *d*).

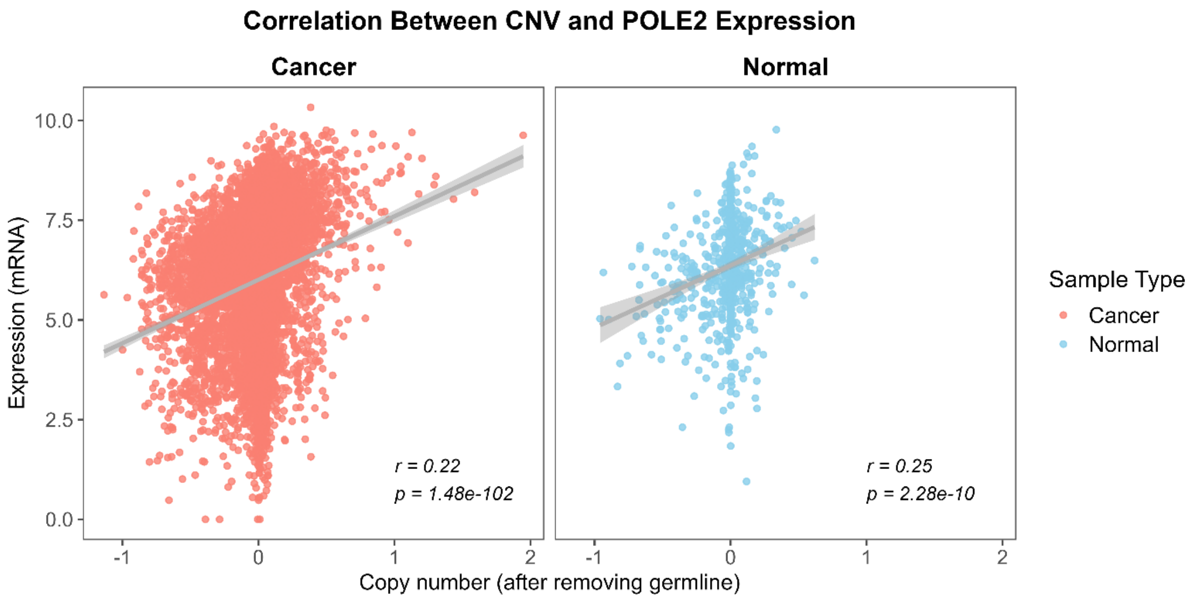


**Figure S3.** Correlation between POLE2 expression and copy number variation (CNV) in cancer and normal samples. Data were obtained from the UCSC Xena platform.

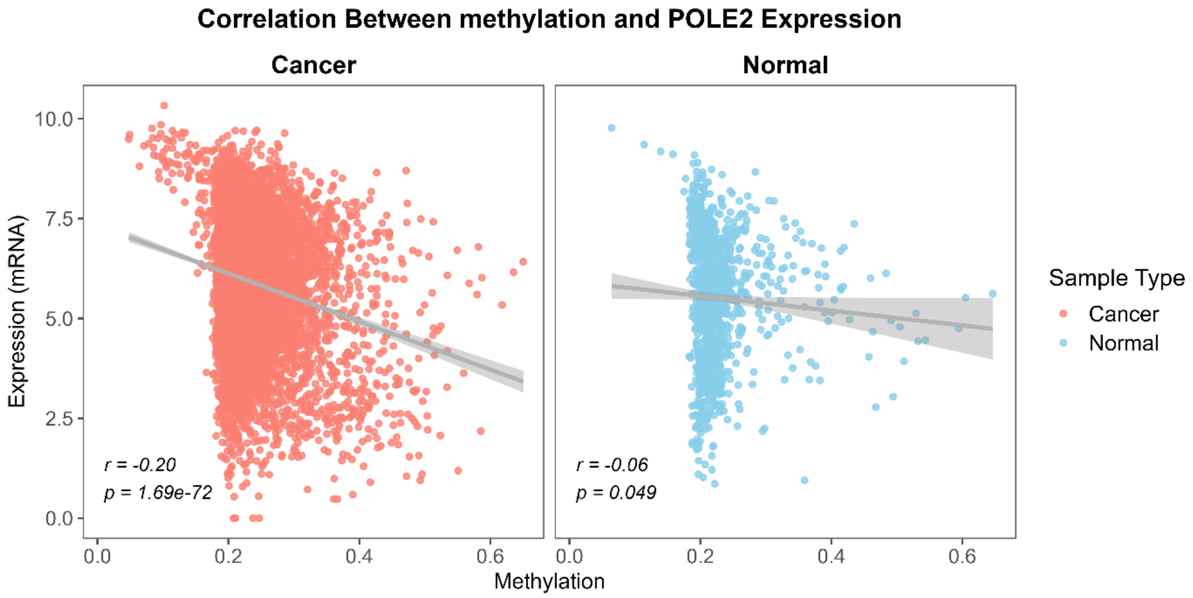


**Figure S4. .** Correlation between POLE2 expression and methylation level in cancer and normal samples. Data were obtained from the UCSC Xena platform.

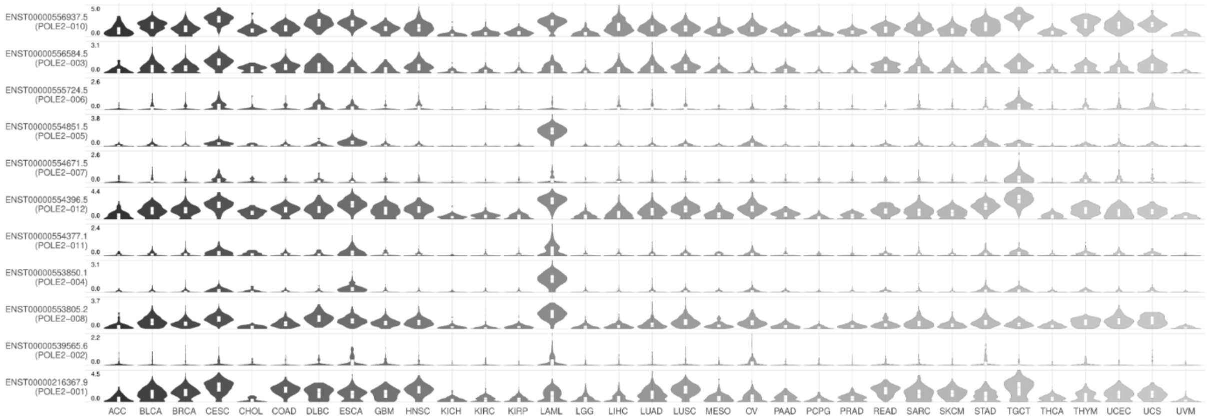


**Figure S5. POLE2 isoforms expression in TCGA cancers**

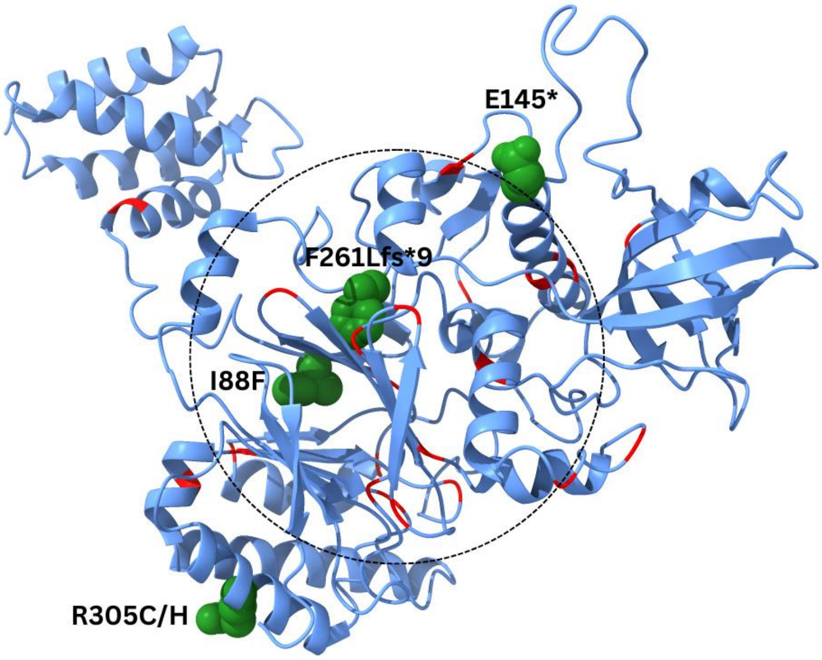


**Figure S6. POLE2 structure with COSMIC-annotated mutations.** Structural representation of the POLE2 protein based on the Protein Data Bank (PDB) entry 7PFO. COSMIC-annotated mutations are indicated in red. Mutations with a recurrence count greater than four are highlighted and labeled in green. The dotted circle denotes a region with a higher occurrence of mutations compared to other parts of the protein.

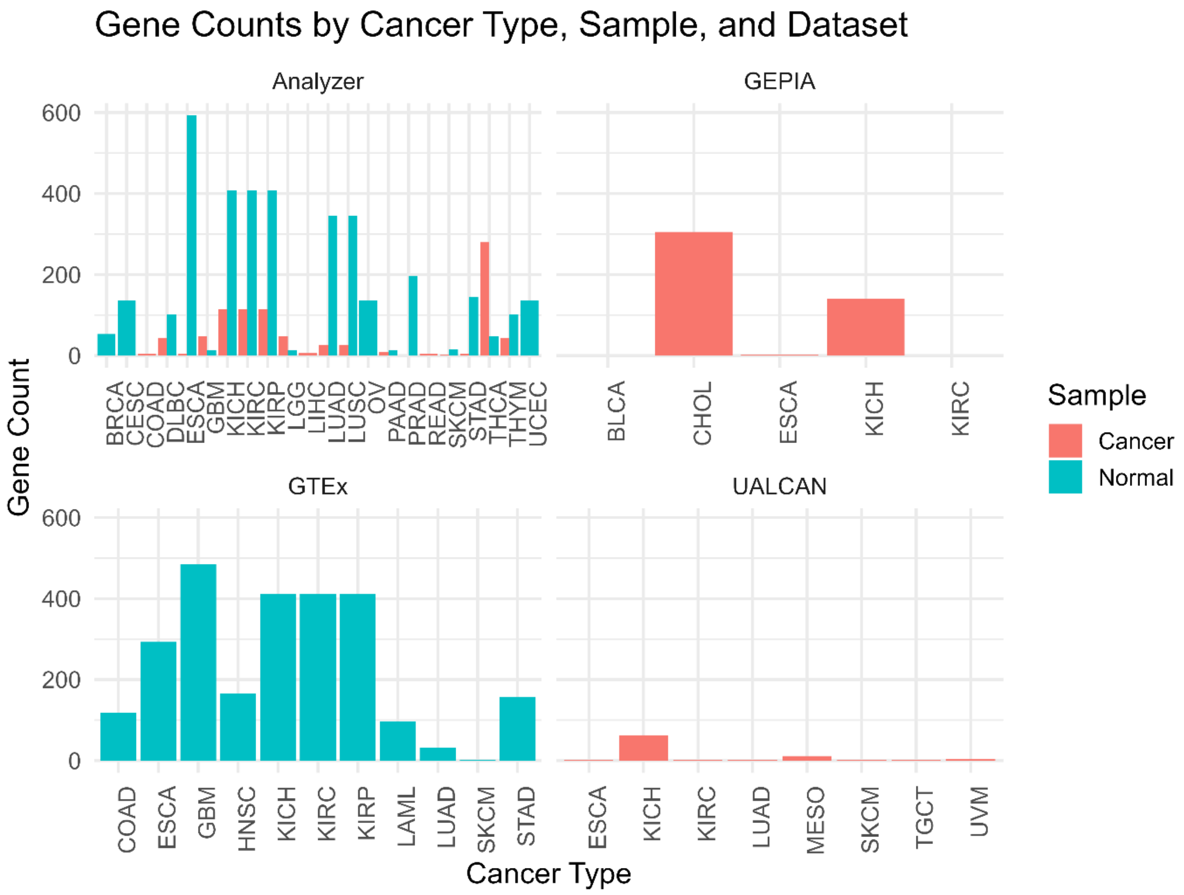


**Figure S7. Number of genes strongly correlated with POLE2 (PCC > 0.8) across multiple datasets**

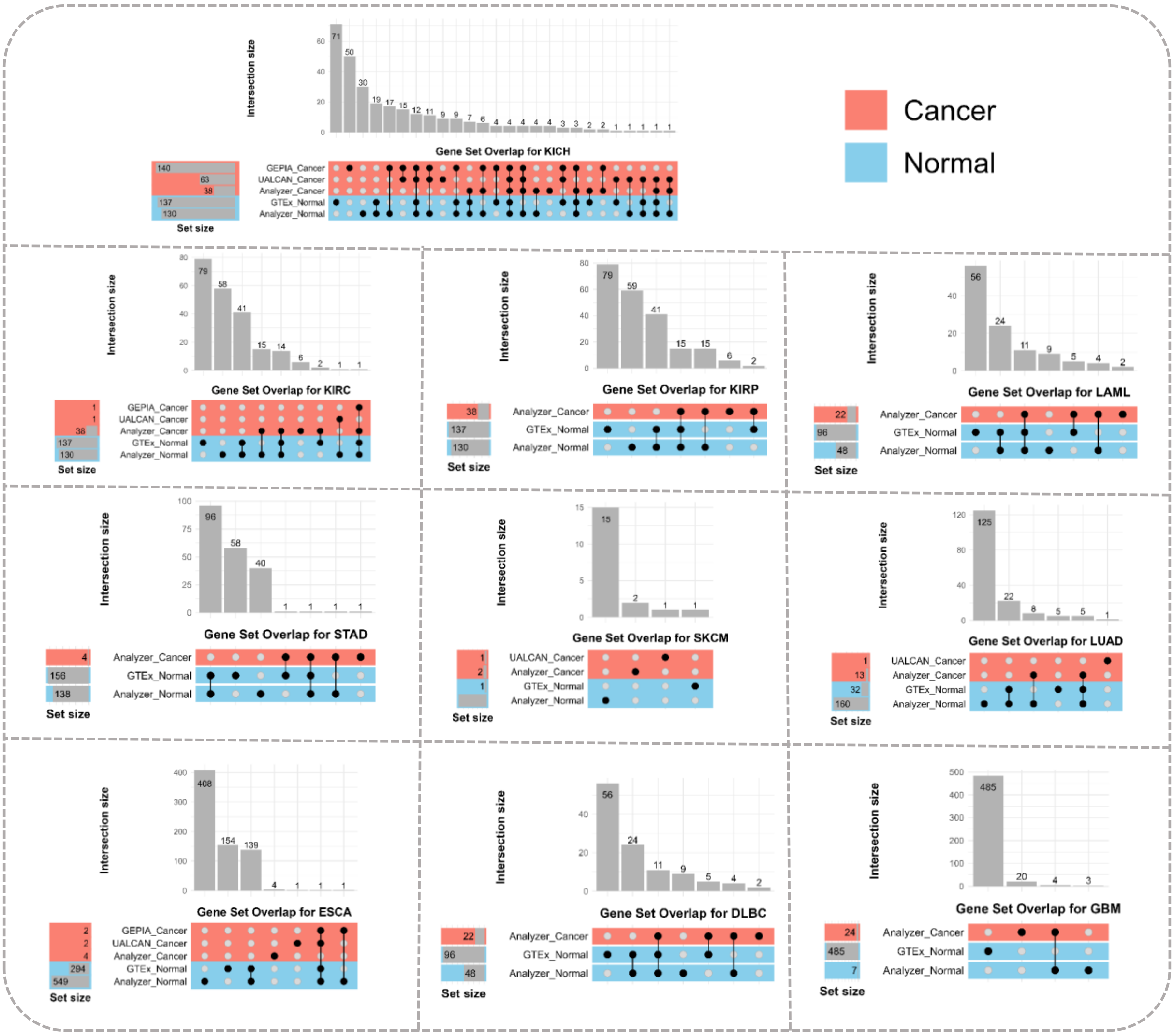


**Figure S8. Overlap of genes highly correlated with POLE2 across multiple datasets in cancer and normal tissues for separate TCGA cancers.**

UpSet plot showing the intersection of genes with a Pearson correlation coefficient > 0.8 with POLE2 expression, derived from six datasets from 4 independent databases: GTEx, GEPIA, UALCAN, and AnalyzeR, each separated by sample type. Top bars indicate the number of overlapping genes shared among dataset combinations, while the left bars show the total number of highly correlated genes per dataset.

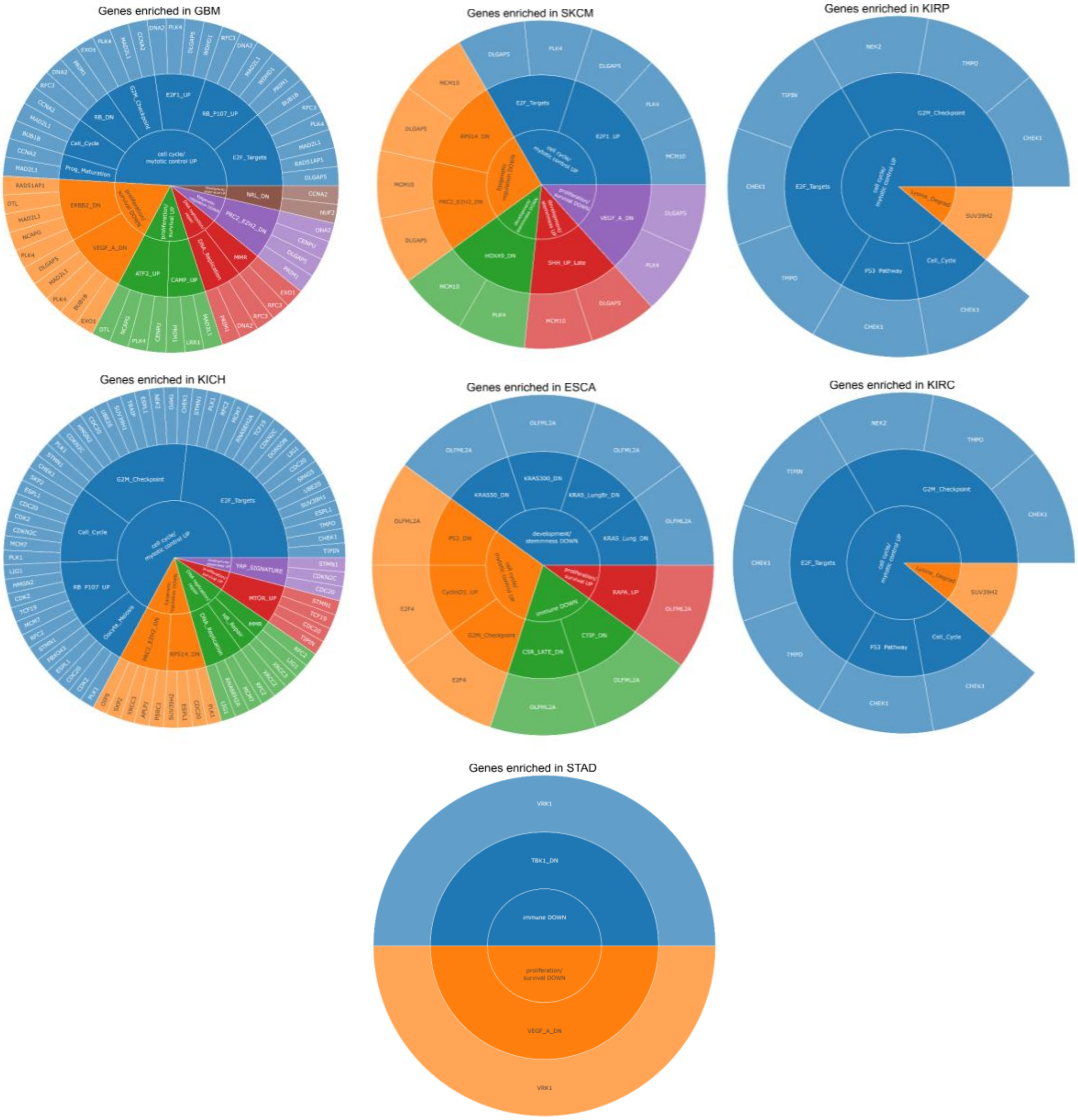


**Figure S9. Sunburst plot of genes from enriched categories correlated with POLE2 in cancer-specific analysis.**
Functionally enriched gene categories, with gene groupings organized hierarchically based on their biological relevance and correlation with POLE2 expression across cancer types.
